# The Role of Alpha-Synuclein in Synucleinopathy: Impact on Lipid Regulation at Mitochondria–ER Membranes

**DOI:** 10.1101/2024.06.17.599406

**Authors:** Peter A. Barbuti, Cristina Guardia-Laguarta, Taekyung Yun, Zena K. Chatila, Xena. Flowers, Bruno FR. Santos, Simone B. Larsen, Nobutaka Hattori, Elizabeth Bradshaw, Ulf Dettmer, Saranna Fanning, Manon Vilas, Hasini Reddy, Andrew F. Teich, Rejko Krüger, Estela Area-Gomez, Serge Przedborski

## Abstract

The protein alpha-synuclein (αSyn) plays a critical role in the pathogenesis of synucleinopathy, which includes Parkinson’s disease and multiple system atrophy, and mounting evidence suggests that lipid dyshomeostasis is a critical phenotype in these neurodegenerative conditions. Previously, we identified that αSyn localizes to mitochondria-associated endoplasmic reticulum membranes (MAMs), temporary functional domains containing proteins that regulate lipid metabolism, including the de novo synthesis of phosphatidylserine. In the present study, we have analyzed the lipid composition of postmortem human samples, focusing on the substantia nigra pars compacta of Parkinson’s disease and controls, as well as three less affected brain regions of Parkinson’s donors. To further assess synucleinopathy-related lipidome alterations, similar analyses were performed on the striatum of multiple system atrophy cases. Our data show region-and disease-specific changes in the levels of lipid species. Specifically, our data revealed alterations in the levels of specific phosphatidylserine species in brain areas most affected in Parkinson’s disease. Some of these alterations, albeit to a lesser degree, are also observed multiples system atrophy. Using induced pluripotent stem cell-derived neurons, we show that αSyn contributes to regulating phosphatidylserine metabolism at MAM domains, and that αSyn dosage parallels the perturbation in phosphatidylserine levels. Our results support the notion that αSyn pathophysiology is linked to the dysregulation of lipid homeostasis, which may contribute to the vulnerability of specific brain regions in synucleinopathy. These findings have significant therapeutic implications.

**Significance Statement:** Synucleinopathy is a complex group of neurodegenerative disorders whose causes and underlying mechanisms remain unknown. In this work, we examined synucleinopathy postmortem brain samples and patient-derived neuron models and identified the functional impairment of the mitochondrial-associated endoplasmic reticulum membrane (MAM) domain, which facilitates lipid regulation. The protein alpha-synuclein is associated with synucleinopathy and increasing levels result in the mislocalization of this protein and the disruption of MAM domains, which, in turn, results in lipid and membrane composition alterations. Specifically, we report that increased alpha-synuclein expression impairs the regulation of phosphatidylserine synthase 2 and the levels of phosphatidylserine in cellular membranes from affected cells. Our study offers mechanistic insight tying alpha-synuclein pathology and lipid dysregulation as seminal factors in synucleinopathy, which may have pathogenic and therapeutic implications.

## Introduction

Synucleinopathy refers to a cluster of adult-onset neurodegenerative conditions marked by a buildup of alpha-synuclein (αSyn) protein aggregates within neuronal cell bodies and fibers, and occasionally within glial cells (1). These disorders primarily encompass Parkinson’s disease (PD), dementia with Lewy bodies (DLB), and multiple system atrophy (MSA) (1). They manifest in various motor and cognitive impairments for which there is currently no cure and a lack of effective disease-modifying treatments. Besides αSyn being used as a biomarker for diagnosis of synucleinopathy (2, 3), a host of genetic studies have provided evidence of its role in synucleinopathy-related neurodegeneration (4). Polymorphisms in the gene encoding αSyn, *SNCA,* that result in increased αSyn expression, have been associated with sporadic PD development (5, 6), and genomic *SNCA* multiplication and rare dominantly inherited *SNCA* point mutations (e.g., A30P and A53T) with familial PD (4). The duplication or triplication of the *SNCA* locus results in αSyn expression at 1.5× or 2× the level observed for a single copy of the wild-type *SNCA* locus, associated with a dose-dependent gain of toxic function, such that patients harboring *SNCA* triplication exhibit earlier PD onset, more aggressive clinical severity, and faster disease progression (7).

Despite the genetic evidence mentioned above, the precise mechanisms underlying how αSyn contributes to neuronal dysfunction and death in synucleinopathy remain elusive. The resemblance of αSyn to lipid-binding proteins has long been recognized (8), and subsequent investigations have confirmed its ability to bind lipids, especially phospholipids and fatty acids (9). While most of this highly expressed brain protein is cytosolic, a fraction of αSyn is bound to various lipid membranes, including synaptic vesicles and plasma membranes (9). As highlighted by Musteikyté et al. (10), the interaction between αSyn and lipid membranes is considered crucial for its biological function, and to play a significant role in the abnormal processes associated with αSyn aggregation and toxicity. Changes in membrane physical properties and chemical composition promote the aggregation of αSyn into toxic amyloid fibrils, while aggregated αSyn species bind to lipid membranes, compromising their integrity (10). However, the precise mechanisms underlying this bidirectional neurotoxic scenario of αSyn-lipid membrane interaction have not yet been fully elucidated.

Our published data indicate that αSyn is also recruited to membrane domains localized in the endoplasmic reticulum (ER) in close apposition to mitochondria (11), known as mitochondrial-associated ER membranes (MAMs). We believe this observation is highly relevant to the question of αSyn-lipid membrane crosstalk, as MAMs are transient lipid raft domains within the ER where key regulatory lipid enzymes converge to control membrane homeostasis (12). Supporting this view, we found that cell lines stably expressing αSyn mutations display alterations in their lipidome (13) and in MAM-related lipid metabolism pathways (11).

To expand our work on αSyn and lipid metabolism in synucleinopathy and MAM, we began the present study by defining the lipidomic profiles of selected postmortem brain regions of PD patients. These analyses showed three main changes that differentiate the lipid profile of the severely affected brain region substantia nigra pars compacta (SNpc) in PD patients from both less affected brain regions in PD and the unaffected SNpc in non-PD controls: (i) elevated levels of cholesterol esters (CEs), (ii) reduced levels of specific ceramide (Cer) species, and (iii) the presence of specific phosphatidylcholine (PC) and phosphatidylserine (PS) species bound to long-chain and unsaturated fatty acids (FAs) that were not present in other PD brain regions or control SNpc. These findings suggest that lipid metabolism changes in PD are specific to certain regions, with the SNpc displaying unique lipid profiles compared to other brain regions affected by PD and control samples. To determine whether the observed lipid changes are specific to PD, we perform lipidomic analysis in MSA as an example of another synucleinopathy. These additional analyses revealed similarities in lipid alterations between MSA and PD samples. However, the changes in lipid species were fewer and of lesser magnitude in MSA compared to PD, suggesting that several lipid alterations appear to be shared by PD and MSA, while others may be specific to vulnerable areas in PD.

In the second part of this study, we turned our attention to a simplified system provided by human induced pluripotent stem cell (iPSC)-derived midbrain neurons to elucidate the cell-autonomous mechanisms leading to the lipid alterations identified in postmortem tissues. These investigations revealed that iPSC-derived neurons harboring *SNCA* duplication recapitulated some of the lipidome changes observed in SNpc samples from PD cases. They also indicated that αSyn plays a physiological role in the modulation of lipid enzymes through its localization to MAM and that increased levels of αSyn interfere with this function, leading to defects in lipid homeostasis. Our findings indicate that αSyn pathology is associated with disrupted lipid homeostasis, potentially contributing to the susceptibility of specific brain regions in synucleinopathy. These insights carry important therapeutic implications.

## Results

### Altered lipidome in the postmortem PD brain

Studies conducted by us and others have reported alterations in lipid profiles in biofluids obtained from individuals with synucleinopathy compared with those from control individuals (13–18). However, the molecular mechanisms underlying lipid changes associated with αSyn pathology and the differential susceptibility of brain regions in synucleinopathy remain unclear. In the present study, we aimed to address these questions by conducting an unbiased lipidomics analysis of postmortem samples from different brain regions obtained from 16 patients with PD and 14 aged-matched control individuals (**Supplementary Table 1**). Because the SNpc has been identified as a primary site of neuropathological changes in PD (19), we specifically compared the lipid profiles of the SNpc between patients with PD and non-PD controls. In addition to the SNpc, we also analyzed three other brain regions from the same cohort of PD donors, the ventral tegmental area (VTA), the substantia innominata (SI), and the rostral hypothalamus (Hypo). These regions provide a range of neurodegeneration, with the greatest degree of neuronal loss in the SNpc, followed by the VTA, the SI, and the lowest degree of neuronal loss in the rostral Hypo (20–25). All collected samples were promptly flash-frozen in the presence of antioxidants (butylated hydroxytoluene) and subsequently processed for lipidomics analysis using ultra-high performance liquid chromatography coupled with tandem mass spectrometry, as previously described (13). Lipids were extracted from 100 µg of tissue from each sample via the chloroform–methanol extraction, followed by a modified Bligh and Dyer protocol.

Triplicate aliquots were analyzed for each sample, allowing us to detect more than 500 lipid species belonging to 31 different classes (**Supplementary Table 2**). Using spiked internal standards with known concentrations, we determined the concentration of each lipid species. We applied the normalization using the optimal selection of multiple internal standards method (26) and conducted the Principal Component Analysis (PCA) to identify samples outside of the 95% confidence interval, which we classified as outliers (**Supplementary Fig. 1**). Subsequently, we compared the lipidome results from all brain areas under study by one way ANOVA and plotted p values for each species in a 3D volcano graph. The data for subset groups were reduced to a 2D polar coordinate system, as explained (27) (**Supplementary Fig. 2A**). The three-way volcano plots helped us visualize that the groups display significant differences in their lipid composition. In particular, our data revealed some significant changes in the levels of specific lipids of cholesteryl esters (CEs) and diacylglycerides (DGs) especially those bound to oleic acid (CE 18.1 and DG 38.1), as well as various phospholipids such as those PC species bound to saturated fatty acids (PC 32.0) and those containing arachidonic acid (PC 38.4) (**Supplementary Fig. 2A**).

To validate these results, we next employed a random forest (RF) machine learning classifier to identify those lipid species with the strongest ability to discriminate among the various conditions, as previously described (16). This approach revealed several lipid classes and species capable of discriminating between PD and control SNpc as well as SNpc and other brain areas (VTA, Hypo and SI) from the same PD cases (**Table 1**). The RF analysis identified that alterations in CEs, (CE 18:1, CE 20:3 and CE 22:6) and phosphatidylinositol (PI) and phosphatidylethanolamine plasmalogen (PEp) species with polyunsaturated acyl chains (PI 40:5 and PEp 36:4), are critical discriminatory variables among brain regions from the same cases and control group.

**Table 1:**
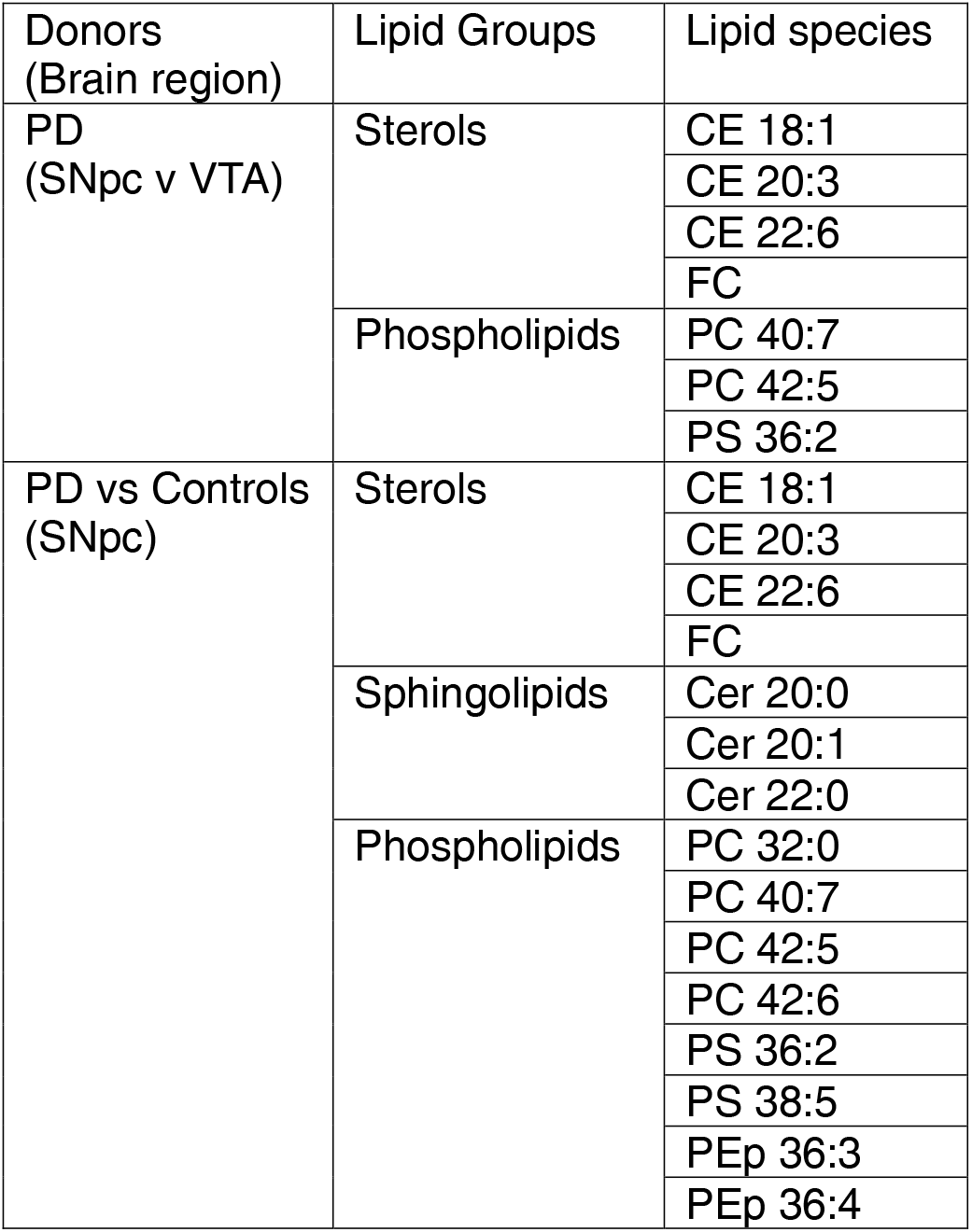
Lipid alterations in postmortem PD brains. Lipid species identified by random forest (RF) classification can discriminate within PD brains differentially susceptible brain regions (substantia nigra pars compacta (SNpc) vs ventral tegmental area (VTA) (upper row) and between PD brains and control brains at the SNpc (lower row). Abbreviations: SNpc, substantia nigra pars compacta; VTA, ventral tegmental area; CE, cholesterol esters; FC, free (unesterified) cholesterol; PC, phosphatidylcholine; PS, phosphatidylserine; Cer, ceramide; PEp, plasmalogen phosphatidylethanolamine.

Other polyunsaturated phospholipids belonging to PE and PEp classes (PEp 36:3, PEp 36:5, PEp 38:3, PEp 38:2, PE 36:2, PE 38:2) were also found to be specifically altered in SNpc from PD, but only when compared to controls. Additionally, two Cer species bound to C22 fatty acids (Cer 22:0 and Cer 22:1) were classified as discriminatory variables able to distinguish between PD vs. control SNpc.

Minimal depth of the RF algorithm shows the distance between the root and the decision nodes using the particular lipid species in the decision trees for the classification. Higher frequencies at shorter distance would indicate that some lipid species are more effective at classifying the different groups. Our results suggest that, although at lower frequencies, defects in the levels of free cholesterol and diglyceride (DG) species, such as DG 38:1 or DG 38:3, are phenotypes unique to PD SNpc compared to controls (**Supplementary Fig. 2B**).

To validate these data, we also compared the levels of various lipid classes and species between PD and control samples and calculated fold-change values. In agreement with our RF results, SNpc samples from patients with PD exhibited higher levels of specific CE species (**Fig. 1A**) than SNpc samples from controls. Our analysis also revealed notable reductions in the levels of various abundant Cer species in the SNpc of patients with PD relative to the levels observed in other areas of the brain and control SNpc, particularly those linked to long-chain FAs (**Fig. 1B**). Furthermore, we found unique alterations in the levels of major phospholipids in PD SNpc compared to the rest of the PD regions under study and SNpc from controls. Namely, our results show that PD SNpc samples displayed higher levels of PC (**Fig. 1C**) and PS (**Fig. 1D**) species bound to long-chain and unsaturated FAs that were not present in the other PD brain regions or control SNpc. Thus, our analyses reveal alterations in three primary classes of lipids: sterols, phospholipids, and sphingolipids. These changes appear region-specific, with the SNpc in PD exhibiting distinct lipid profiles compared to other PD-affected brain regions and control samples.

**Figure 1:**
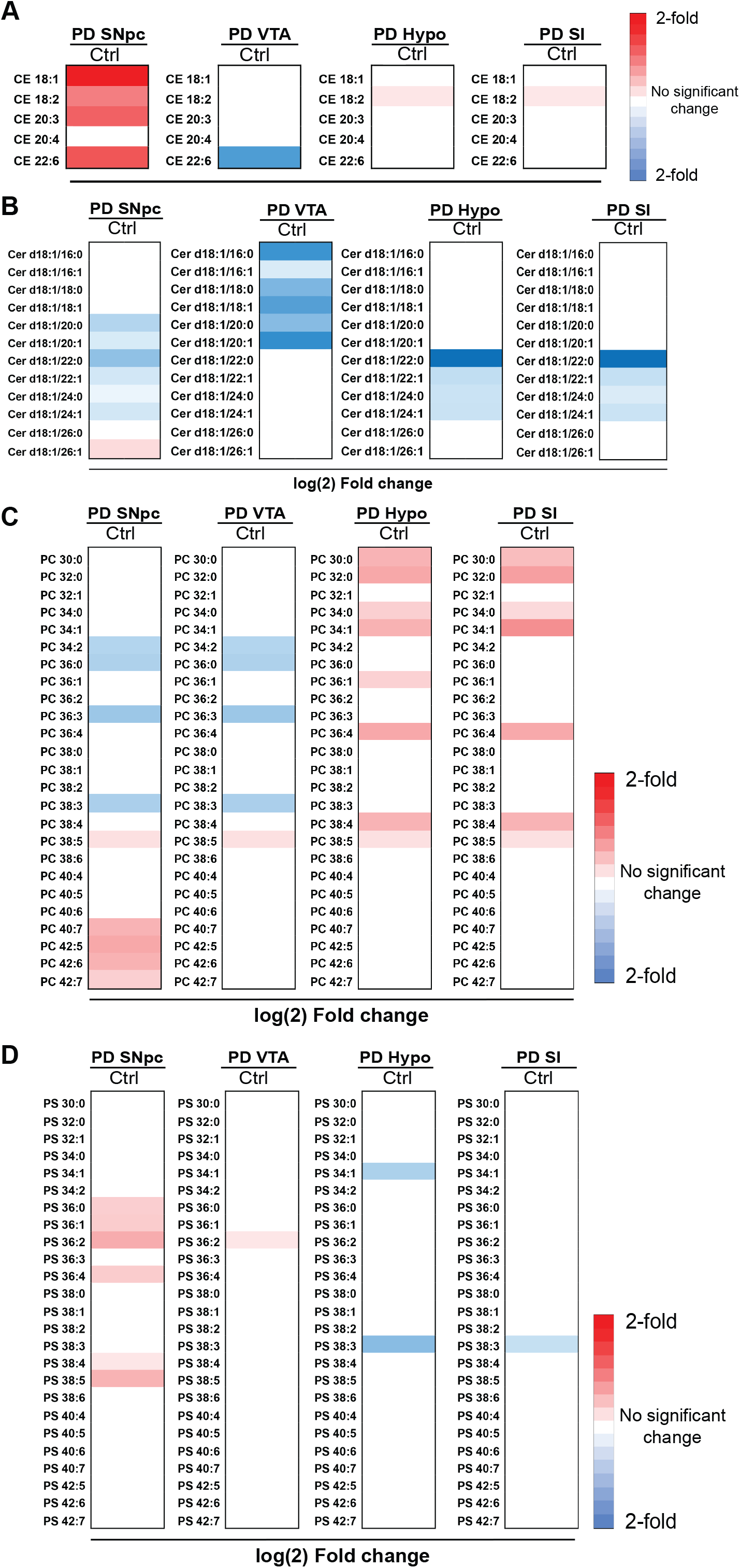
Lipid alterations in the postmortem Parkinson’s disease (PD). (A) Heatmaps representing 2-fold changes in the abundance of individual lipid species of (**A**) esterified cholesterol (CE) (**B**) ceramide (Cer), (**C**) phosphatidylcholine (PC) and (**D**) phosphatidylserine (PS) across different brain regions: substantia nigra pars compacta (SNpc), ventral tegmental area (VTA), hypothalamus (Hypo), substantia innominata (SI) of PD postmortem donors compared to controls.

We next ask whether the lipid changes reported above are specific to PD or shared by other synucleinopathies. Accordingly, we analyzed the lipidome of striatum samples from MSA cases (**Supplementary Table 3**), since it is the MSA primary site of neuropathological changes (19), and compared it to our data from PD and non-PD controls. These analyses revealed that the striatum in MSA has alterations in some of the CE species like those detected in the SNpc of PD cases (**Supplementary Fig. 3A**). In addition, we were also able to find increases in PS species bound to long-chain and unsaturated FAs (e.g., PS 38:4, PS 38:5) like those found in PD SNpc samples. However, the magnitude and extent of the lipid changes in the MSA striata were not as profound as those found in PD SNpc samples and we did not replicate the alterations in free cholesterol, Cer, and PC levels (**Supplementary Fig. 3**). Thus, the partial similarity of lipid alterations between PD SNpc and MSA striatum suggests that some observed changes detected in affected brain areas are shared between these two distinct synucleinopathies. In contrast, others such as alterations in free cholesterol, Cer, and PC seem specific to PD SNpc.

### Lipid alterations in human iPSC-derived neurons are associated with αSyn gene-dosage

To shed light on the mechanisms driving the observed lipid alterations in the brains of patients with synucleinopathy, we used human iPSC-derived neurons as a simplified model system that would be more suitable for biochemical and molecular investigations. Because alterations in αSyn expression are strongly implicated in PD pathogenesis (28), we sought to assess the impact of αSyn dosage on neuronal lipid metabolism.

Thus, cultured iPSC-derived neurons were used to conduct the same lipidomics analyses applied to brain samples using iPSC lines expressing varying levels of αSyn, including iPSCs, in which αSyn expression was knocked out (αSyn-KO), a line expressing endogenous αSyn levels (αSyn-NL), and a line expressing 1.5× endogenous levels (αSyn-Duplication). These human fibroblast-reprogrammed iPSC lines were obtained from different individuals, and their characterizations are detailed in **Supplementary Table 4,** and previous publications (29–33). As previously described, iPSCs were first differentiated into neural precursor cells, which were used to derive human neurons, as reported previously (29–31, 33–35). After 30 days *in vitro* (DIV30) under directed neuronal differentiation conditions, >90% of neural precursor cells differentiated into neurons, as evidenced by the expression of the neuronal markers Tuj1 and Map2. Among these neurons, ∼20%–30% expressed dopaminergic markers, including tyrosine hydroxylase and dopamine transporter, and were characterized by a ventral midbrain identity, as evidenced by the expression of FoxA2, in keeping with previous studies (30, 36) (**Supplementary Fig. 4A–C).** Using immunoblot analysis, we confirmed that iPSC-derived αSyn-Duplication neurons expressed 50% more αSyn than age-matched αSyn-NL neurons, and that iPSC-derived αSyn-KO neurons did not express αSyn (**Supplementary Fig. 4D**) (35, 37). No differences were observed in differentiation efficiencies or survival rates among the different iPSC-derived neuronal lines.

While our lipidomic analysis of αSyn-Duplication neurons did not detect the kind of alterations in sterols observed in PD whole tissue SNpc extracts, we found substantial changes in sphingolipids in these neurons compared to both αSyn-NL and αSyn-KO neurons. Indeed, we found increases in the overall levels of Cer and its saturated precursor dihydro-Cer (**Fig. 2**), as well as in those of the dihydrosphingomyelin (dhSM) lipid class. However, when all the species from these sphingolipid classes were analyzed, we observed, like in PD SNpc, a marked decrease in those containing long-chain FAs in αSyn-Duplication neurons compared to controls (**Fig. 2A**). More, complex sphingolipids such as monohexosyl-Cer and lacto-Cer were reduced in both overall abundances and in the abundance of species containing long-chain FAs in αSyn-Duplication neurons compared with control neurons, whereas αSyn-KO neurons exhibited increases in the levels of these species (**Fig. 2A**, **Supplementary Fig. 4E**); a similar observation was made for the ganglioside mono-sialodihexosylganglioside (**Fig. 2A**, **Supplementary Fig. 4E**). Concomitantly, αSyn-Duplication neurons displayed increases in sphingolipid species with shorter acyl chains, while αSyn-KO neurons presented the inverse phenotype (**Fig. 2A** and **Supplementary Fig. 4F**), implying that these lipid alterations are inversely correlated with the αSyn dosage.

**Figure 2:**
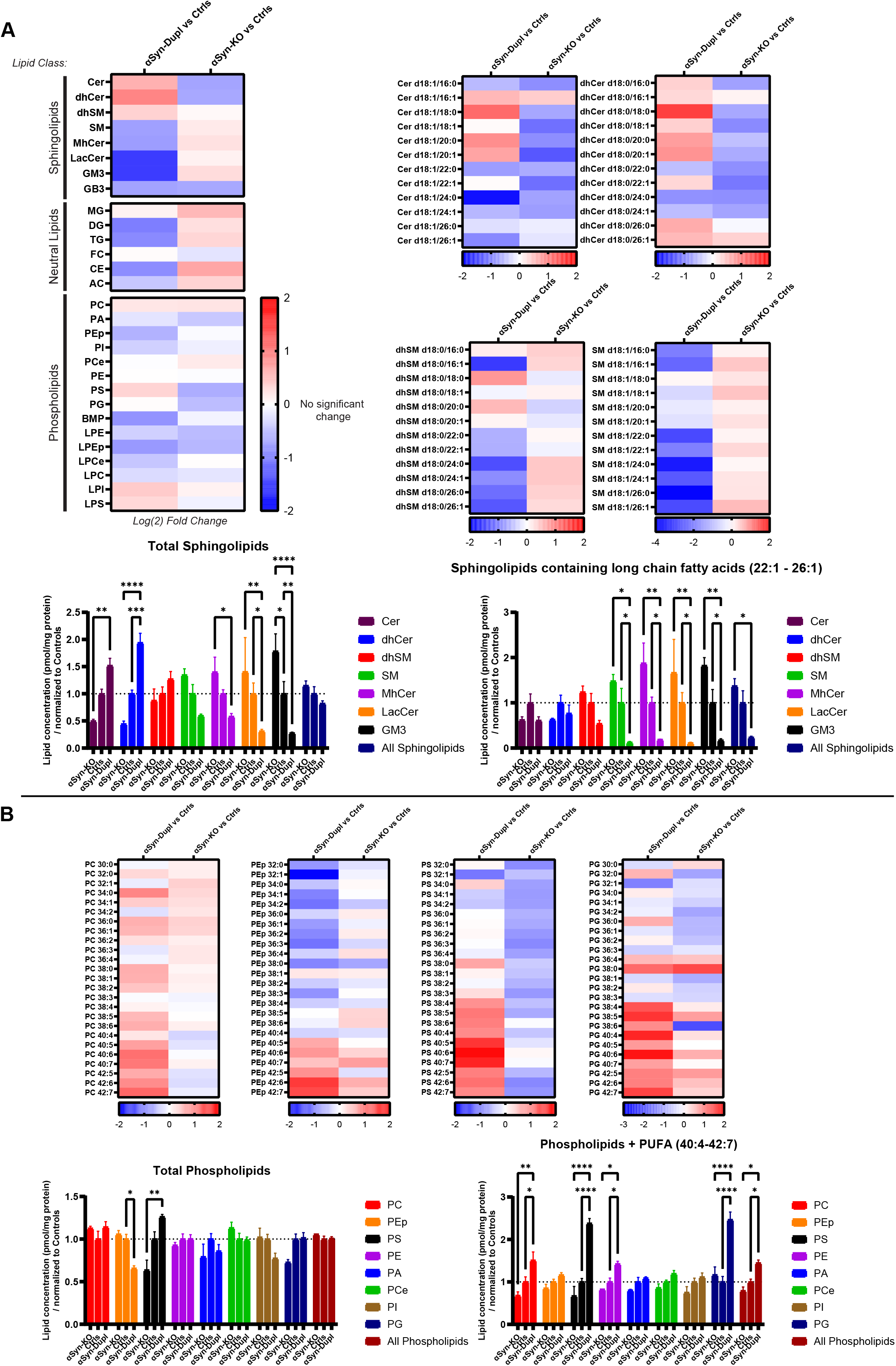
Lipid alterations in patient-derived neurons expressing differing alpha-synuclein (αSyn) doses. (**A**) Lipidomic heat maps showing Log_2_ fold changes in lipid groups and selected individual lipid species in induced pluripotent stem cell (iPSC)-derived neurons carrying either αSyn duplication (αSyn-Dupl) or αSyn knock out (αSyn-KO), normalized to age-matched neurons expressing endogenous αSyn levels (Ctrls) after 30 days of directed differentiation. Relative concentrations of total sphingolipids (**bottom left**) and sphingolipids containing long-chain fatty acids (22:1 to 26:1) (**bottom right**) are presented, normalized to the concentrations in Ctrl samples. Data are presented as the mean ± SEM of at least 3 independent biological replicates (n) analyzed by an ordinary two-way analysis of variance (ANOVA), followed by Tukey’s multiple comparison test. Total sphingolipids: Interaction (F_16,99_ = 5.462; ****p<0.0001) and Row Factor (cell line) (F_2,99_ = 1.743; p=0.1804). Sphingolipids containing long-chain fatty acids: Interaction (F_16,99_ = 1.522; p=0.1068) and Row Factor (cell line) (F_2,99_ = 19.09; ****p<0.0001). (**B, top**) Lipidomic heat maps displaying Log_2_ fold changes in selected phospholipids. (**Bottom**) Relative concentrations of total phospholipids and phospholipids containing polyunsaturated fatty acids (PUFAs, 40:4 to 42:7) are presented, normalized to the concentrations in Ctrl samples. Data are presented as the mean ± SEM of at least 3 independent biological replicates (n) analyzed by an ordinary two-way ANOVA, followed by Tukey’s multiple comparison test. Total phospholipids: Interaction (F_18,110_ = 2.485; **p=0.002) and Row Factor (cell line) (F_2,110_ = 1.684; p=0.1904). Sphingolipids containing long-chain fatty acids: Interaction (F_18,110_ = 5.215; ****p<0.0001) and Row Factor (cell line) (F_2,110_ = 67.56; ****p<0.0001).

As for phospholipids, consistent with our findings in PD SNpc, αSyn-Duplication neurons also displayed marked differences in this lipid class compared with control and αSyn-KO neurons. Notably, we detected an increase in the concentration of PS as well as all species of phospholipids bound to polyunsaturated fatty acids (PUFAs; 40:4 to 42:7), including not only PS and PC but also phosphatidylethanolamine (PE) (**Fig. 2B** and **Supplementary Fig. 4F**). Deviations in the opposite direction were observed for these species in αSyn-KO neurons, suggesting a potential link between the levels of these lipid species and the αSyn dose. Of note, not all phospholipid species were increased as the concentration of PEp was decreased in αSyn-Duplication neurons compared with αSyn-NL neurons. Thus, our whole-cell lipidomic analysis of αSyn-duplication neurons, while not detecting changes in sterols as observed in PD SNpc, provided a more detailed understanding of the alterations in both sphingolipids and phospholipids in connection to αSyn dosage. This underscores the value of our cell model in exploring the association between αSyn and lipid metabolism changes in synucleinopathy.

### Analysis of subcellular fractions from cell models expressing endogenous αSyn

The convergence of lipid metabolic pathways on MAM domains regulates cellular lipid homeostasis. Our published data indicate that αSyn can localize to MAM domains and that point mutations in *SNCA* or overexpression of wild-type αSyn disturb the regulation of MAM domains and MAM-associated cellular functions (11). To confirm these findings in iPSC-derived neurons, we first performed subcellular fractionation as in (11) to assess αSyn content of purified ER, MAM, mitochondria, and cytosol by immunoblot. This experiment confirms that αSyn localizes to the MAM and cytosolic fractions in αSyn-NL iPSC-derived neurons, with limited localization in ER fractions (**Fig. 3**). In the total non-fractionated homogenate, the αSyn-Duplication line expresses 1.5× the level of αSyn detected the αSyn-NL line, consistent with the duplication phenotype (**Supplementary Fig. 4D**); however, the relative quantification of subcellular fractions from αSyn-Duplication neurons revealed the relative enrichment of αSyn in the MAM and ER fractions, with levels 3× and 4× those found in the respective fractions from αSyn-NL neurons. These findings indicate that the αSyn localization is altered when the concentration increases above endogenous physiological levels. This leads to the selective accumulation and association of αSyn at MAM and ER domains (**Fig. 3A**).

**Figure 3:**
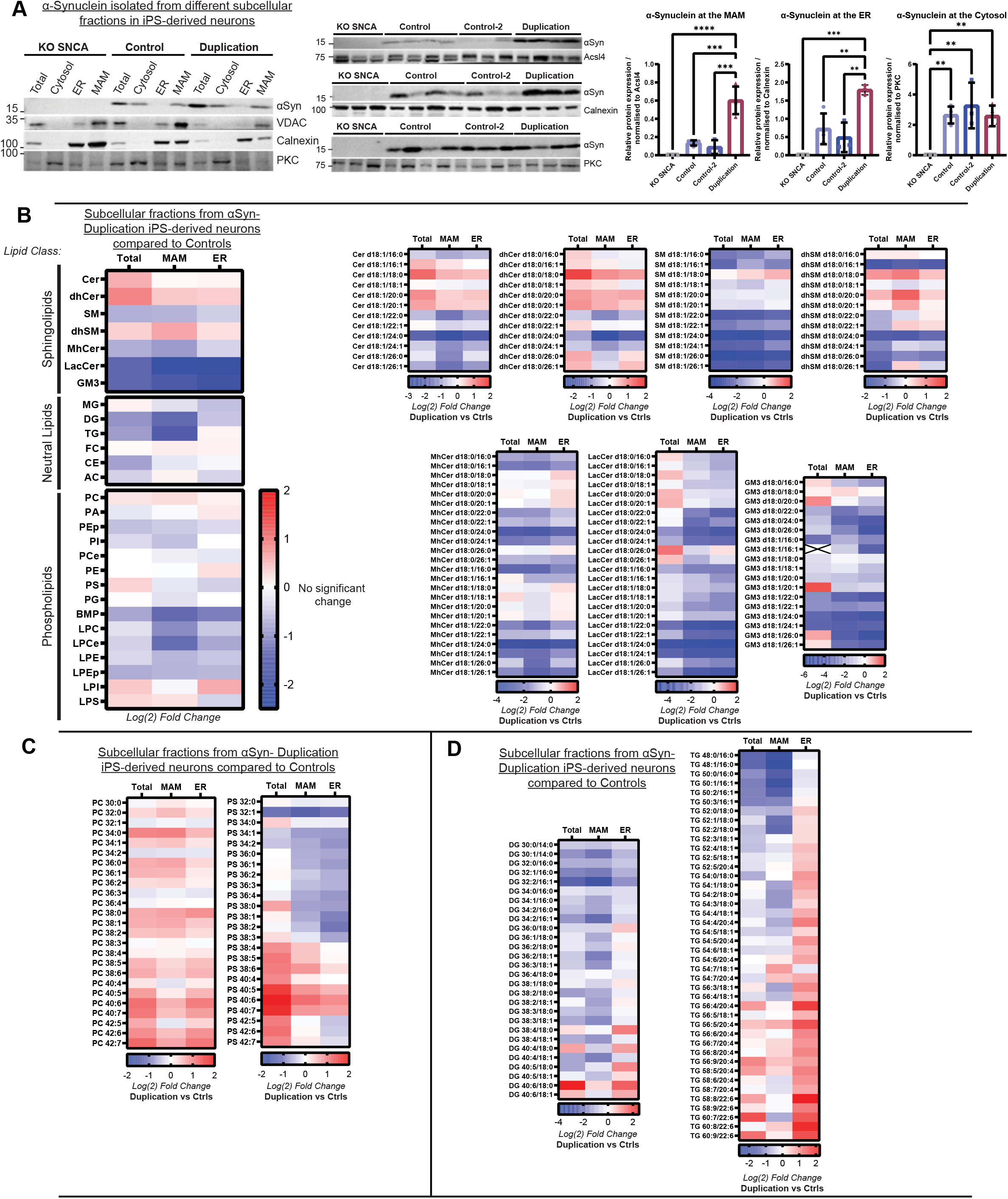
Lipid alterations in neurons with alpha-synuclein (αSyn) duplication across subcellular fractions. (**A**) Qualitative (**left**) and quantitative (**right**) assessments of monomeric αSyn distributions in the total unfractionated homogenate and in mitochondria-associated endoplasmic reticulum (ER) membrane (MAM), ER, and cytosol fractions from induced pluripotent stem cell (iPSC)-derived neurons carrying different αSyn levels after 30 days of directed differentiation. A semi-quantitative assessment of αSyn abundance was performed using protein immunoblots. Protein levels were normalized to the levels of long-chain fatty acid– CoA ligase 4 (Acsl4, MAM marker), calnexin (bulk ER marker), or protein kinase C (PKC, cytosolic marker). Data are presented as the mean ± SD of at least 3 independent biological replicates (n) analyzed by an ordinary one-way analysis of variance (ANOVA). MAM (Interaction F_3,10_ =32.56 ****p<0.0001); ER (Interaction F_3,9_ =17.21 ***p=0.0005); Cytosol (Interaction F_3,10_ =9.424 **p=0.0029). (**B**) Lipidomic heat maps displaying Log_2_ fold changes in groups and individual lipid species of sphingolipids; neutral lipids, and phospholipids in αSyn-Duplication iPSC-derived neurons compared with controls (Ctrls) in the total unfractionated homogenate and in MAM and ER fractions. In Ctrl neurons, alterations in sphingolipid species were consistent between MAM and ER fractions (**right**). (**C**) MAM and ER fractions show elevated phosphatidylcholine and phosphatidylserine concentrations, but a specific reduction in phosphatidylserine lipid species comprised of smaller-chained hydrocarbons is also observed. (**D**) Individual neutral lipids species across the total homogenate, MAM and ER fractions highlights notable differences in diglyceride (DG) and triglyceride (TG) lipids between fractions.

The MAM domain is a transient lipid raft induced by the clustering of cholesterol, sphingomyelin (SM), and saturated phospholipids (38) that modulates specific protein subsets (39). Alterations in the lipid compositions of MAM domains impair the enzymatic activities that localize and are regulated in these regions (40, 41). Thus, next, we applied our lipidomics analysis to MAM and ER fractions from αSyn-Duplication and αSyn-NL neurons to expand our analysis of lipid compositions in these cell lines. Consistent with our whole cell lipidomic analysis, our data did not reveal any alterations in sterols in MAM and ER fractions among genotypes but both in sphingolipids and phospholipids. Indeed, in the case of sphingolipids, we found that the levels of SM and complex sphingolipids were reduced in αSyn-Duplication neurons compared with αSyn-NL neurons, and a reciprocal relationship between Cer and SM was noted in the total homogenates and in ER and MAM fractions (**Fig. 3B, Supplementary Fig. 5A**). Concomitantly, the overall concentration of dihydro-SM species increased in MAM fractions from αSyn-Duplication neurons compared with MAM fractions from αSyn-NL neurons (**Fig. 3B**). Further lipidomics analysis indicated an imbalance in sphingolipid species in αSyn-Duplication neurons compared with αSyn-NL neurons, characterized by a marked decrease in species bound to long acyl chains and a slight increase in shorter and saturated species from the dihydro-SM, dihydro-Cer, and Cer classes (**Fig. 3B)**. Interestingly, subcellular fractions from αSyn-KO neurons showed the opposite phenotype of those from αSyn-Duplication neurons, further supporting the potential contribution of αSyn levels to the modulation of sphingolipid homeostasis (**Supplementary Fig. 5A-B**).

Regarding phospholipids, similar to the alterations observed in total homogenates, the MAM and ER fractions of αSyn-Duplication neurons displayed elevated levels of PC and PS species bound to PUFAs compared with those fractions from αSyn-NL neurons (**Fig. 3C**). Because PS is synthesized at MAM domains, the disparity between PS bound to long-chain PUFAs and PS bound to shorter-chain and more saturated fatty acyl chains was particularly striking, suggesting that the metabolism of PS is impaired at MAM domains in αSyn-Duplication neurons (**Fig. 3D**). As before, αSyn-KO neurons showed the inverse phenotype to the phenotype observed in αSyn-Duplication neurons, implying a positive relationship between αSyn dosage and PS metabolism (**Supplementary Fig. 5C**).

Lipid changes we did not observe in either whole SNpc tissue from PD or iPSC neuron extracts from αSyn-Duplication neurons were the high concentrations of certain diglyceride and triglyceride species in MAM and ER fractions (**Fig. 3D**). These elevations were particularly striking in triacylglycerol (TG) species containing oleic acid (C18:1) and long PUFAs, such as arachidonic acid (C20:4) and docosahexaenoic acid (DHA) (C22:6). As such, they introduce the intriguing possibility that the *de novo* synthesis of TG occurs in the ER of αSyn-Duplication neurons.

Altogether, our lipidomics results reveal an association between high levels of αSyn and the disruption of membrane composition and the lipid milieu of ER and MAM fractions, which can help explain previously reported defects in MAM activities and ER–mitochondria crosstalk (11).

### Alterations in the PS synthetic machinery parallel αSyn dosage in iPSC-derived neurons

We then examined the impact of αSyn dosage on MAM functionality, focusing on PS metabolism due to the aforementioned alterations in this lipid class. Accordingly, we monitored [^3^H]PS formation and its decarboxylation to [^3^H]PE by incubating our cell models with [^3^H]serine, as previously described (11). Our findings indicated that upon incubation with [^3^H]serine, αSyn-Duplication neurons have a ∼2-fold higher content of [^3^H]PS compared with αSyn-NL neurons and a ∼4-fold higher content compared with αSyn-KO neurons (**Fig. 4A**). Conversely, upon incubation with [^3^H]serine, we found a negative relationship between αSyn dosage and [^3^H]PE content in the same cell models (**Fig. 4A**). These results suggest that αSyn-Duplication is associated with an impaired decarboxylation of PS into PE, albeit we cannot exclude that PS synthesis is also increased in these neurons.

**Figure 4:**
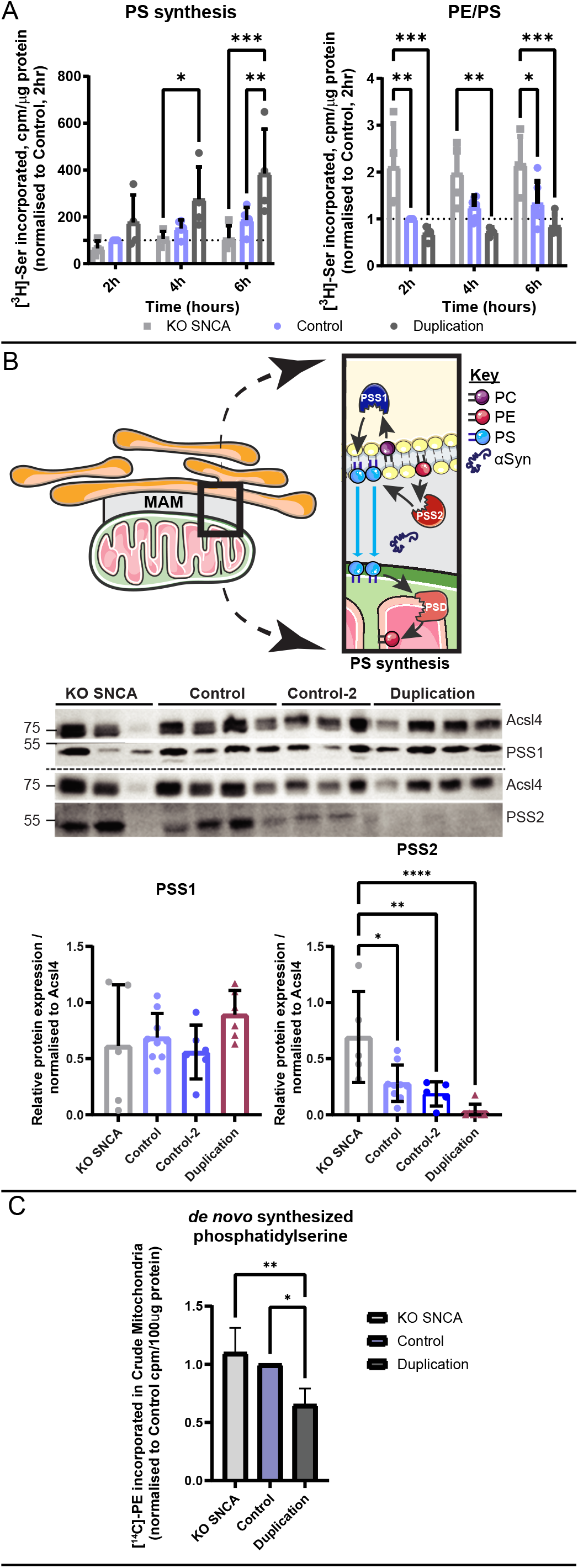
Different alpha-synuclein (αSyn) levels alter mitochondria-associated endoplasmic reticulum membrane (MAM) function. **(A)** Phospholipid synthesis in induced pluripotent stem cell (iPSC)-derived neurons. Quantification of *de novo* phosphatidylserine (PS) synthesis and the ratio phosphatidylethanolamine (PE) to PS (PE/PS) levels in patient-derived neurons after 30 days of directed differentiation after incubation with ^3^H-Serine (^3^H-Ser) for the indicated times. Data were normalized to the Control cell line, and at least 4 independent differentiations were performed. KO SNCA, *SNCA* knock out. A repeated measures two-way analysis of variance (ANOVA) with assumed sphericity was performed for the indicated experiments: PS synthesis: Column (Cell line) factor (F_2,10_ =6.777; *p=0.0138); Row (time) factor (F_2,20=_ 10.92; ***p=0.0006); PE/PS: Cell line factor (F_2,11_ = 13.41; **p=0.0011); time factor (F_2,22=_ 1.180; p=0.3260). **(B**) Protein levels of PS synthase (PSS)1 and PSS2 at the MAM domain. Analysis of MAM fractions from iPSC-derived neurons probed for PSS1 and PSS2, with a representative image shown, adapted from (81). Data are presented as the mean ± SD of at least 4 independent biological replicates (n) analyzed by an ordinary one-way ANOVA of the indicated proteins: PSS1 (Interaction F_3,22_ =1.326; p=0.2912); PSS2: (Interaction F_3,22_ =10.46; ***p=0.0002). Tukey’s multiple comparison was used for post hoc analysis with a single pooled variance. *p<0.05, **p<0.01, ***p<0.001, ****p<0.0001. PC, phosphatidylcholine. **(C)** Assessment of PSS2 activity. Quantification of *de novo* PS synthesis from ^14^C-PE in patient-derived neurons after 30 days of directed differentiation. After subcellular fractionation and quantification, 100 µg of protein isolated from the crude mitochondria was incubated with ^14^C-PE for 45 minutes. Lipids were immediately extracted using the chloroform/methanol extraction method followed by a modified Bligh and Dyer protocol. A minimum of 4 independent differentiations were performed, and data were normalized to the Control cell line. A repeated measure one-way ANOVA with assumed sphericity was performed. Column (Cell line) factor (SS: 0.4430; DF: 2; MS: 0.2215; F_2,6_ =12.23; **p=0.0076).

The MAM domains have been identified as the subcellular locus for PS synthesis by the PS synthase (PSS) enzymes PSS1 and PSS2 (42). Thus, to study further the question of PS metabolism in αSyn-Duplication neurons, we conducted immunoblot analysis to examine the levels of PSS1 and PSS2, which use PC and PE, respectively, as substrates (43) (**Fig. 4B**). MAM fractions isolated from iPSC-derived neurons at DIV30 revealed an inverse relationship between PSS2 immunopositivity and αSyn expression; however, no differences in PSS1 immunopositivity were observed among αSyn-KO, αSyn-NL, and αSyn-Duplication neurons (**Fig. 4B**). Similarly, no differences in PS decarboxylase (PISD) immunopositivity were observed among αSyn-KO, αSyn-NL, and αSyn-Duplication neurons (**Supplementary Fig. 6**). We next used [^14^C]PE to assess PSS2 enzymatic activity in isolated crude mitochondria preparations obtained from αSyn-KO, αSyn-NL, and αSyn-Duplication neurons. In keeping with the immunoblot findings of reduced PSS2 levels in αSyn-Duplication neurons, we observed a reduction in PS synthesis in αSyn-Duplication neurons compared with αSyn-NL and αSyn-KO neurons (**Fig. 4C**). Therefore, taken together, our findings indicate that the dosage of αSyn influences the conversion of PS to PE and the expression of PSS2 through PS negative feedback.

## Discussion

In this study, our objective was to investigate lipid homeostasis in synucleinopathy, starting by examining human post-mortem samples from different brain areas of PD patients and confirming the changes in MSA cases. We then moved to a simplified cell-autonomous model provided by human iPSC-derived neurons expressing varying levels of αSyn to begin our mechanistic investigations into the relationship between αSyn and lipid homeostasis, focusing on MAM.

Our lipid profile analysis using RF classification validated previously identified changes in human clonal cell lines carrying pathogenic αSyn-A53T (13). We observed alterations in CEs, glycerolipids (e.g., TG), glycerophospholipids (e.g., PEp or PC), and sphingolipids (e.g., Cer) that have been consistently observed in patients with PD (44). Some of these changes are also present in other neurodegenerative disorders (45), suggesting a potential commonality in neurodegeneration. Specifically, our RF-classified results highlighted alterations in certain CE species in SNpc from PD cases compared to other PD brain areas and SNpc from non-PD controls. As previously observed by others (46), these CE species were also increased in striatum and cerebellar samples from MSA patients. Similar elevations in CE have been observed in Huntington’s disease (47) and Alzheimer’s disease cases (48), indicating a broader relevance for changes in these lipid species beyond synucleinopathy. Given the increasingly recognized involvement of neuroinflammation in neurodegeneration, including synucleinopathy, it is notable that increases in cholesterol esterification can result from the stimulation of cholesterol traffic and turnover upon microglia activation. These changes allow microglia to adjust their membranes to respond to specific pathogens for phagocytosis or clonal expansion during inflammation. Thus, it is possible that the observed elevations in certain CE species in whole tissue extracts are reporters of pro-inflammatory conditions in those areas most affected in these synucleinopathies. Likewise, SNpc from PD cases and to a lesser degree, striatum samples from MSA patients presented with important elevations in specific species of PS, that were also found in the iPSC-derived neurons carrying the pathogenic αSyn-Duplication.

Relevant to these data, our results confirm that αSyn localizes to MAM domains, where it contributes to the regulation of lipid enzymes, such as PSS1 and PSS2. We posit that αSyn in PD disrupts this function, leading to substantial defects in lipid homeostasis and membrane functionality. Further investigation is required to understand the precise molecular mechanisms governing αSyn’s interactions with these enzymes. However, our lipidomics data from isolated MAM fractions also raises the possibility that αSyn pathology disrupts the reorganization of ER membranes required for MAM formation, which could impair not only PSS1 and PSS2, but also the various enzymatic activities localized to this domain. For example, the change in the lipid milieu after MAM formation induces not only the activation of phospholipid synthesis by enzymes such as PSS1 and PSS2 but also FA acylation by long-chain fatty-acid–CoA ligases (Acsls) (43, 49). These MAM-resident enzymes activate saturated FAs, monosaturated FAs (Acsl1), and PUFAs (Acsl4 and Acsl6), a process vital for maintaining lipid homeostasis and supporting cellular functions related to lipid metabolism. The simultaneous regulation of these enzymes modulates the degree of phospholipid saturation and, thus, membrane fluidity and permeability. Therefore, alterations in the formation and regulation of MAM domains contribute to modifications in phospholipid synthesis, desaturation, and membrane fluidity. In particular, evidence supports the existence of interplay between αSyn and PS. αSyn binds to PS, with a preference for PS comprised of PUFAs (50). The accumulation of insoluble αSyn in PD parallels PS abundance (51) and alterations in *de novo* PS synthesis regulation.

PS plays various roles in diverse biological processes, many of which remain incompletely understood. For example, mammals have evolved to have two biosynthetic pathways for PS generation (52): PSS1 is ubiquitously expressed in all tissues, whereas PSS2 is highly expressed in exclusive brain regions, testis, and kidney. In these tissues, particularly the brain, PS may be so critical for function that an alternative conserved pathway exists as an evolutionary backup; alternatively, a specific requirement and, therefore, function may exist for the PS species generated by each PSS enzyme. Studies have revealed that PSS2, which exclusively uses PE as the substrate (53) for PS generation, preferentially generates high levels of PS featuring reduced lipid chain lengths and containing DHA (54). In contrast, PSS1, which has a preference for PC but does not use it exclusively as a substrate for PS generation, preferentially generates PS rich in PUFAs (e.g., PS 40:6), which are the most abundant PS species in the brain (55) and are likely composed of steric acid (18:0) and DHA. In our study, αSyn levels paralleled the *de novo* synthesis and abundance of PS in iPSC-derived neurons, which, in turn, was associated with reduced PSS2 levels at MAM domains. Studies comparing the activities of PSS enzymes in the brains of patients with PD and healthy controls have revealed reduced PSS enzymatic activity levels in the SNpc comparted to the putamen and cortical regions in control brains (56). However, patients with PD display increased PSS enzymatic activity, specifically in the SNpc (56).

Additionally, the activities of the rate-limiting enzymes involved in PC and PE synthesis (PCCT and PECT, respectively) also increase in the PD SNpc compared with the rates in healthy controls (56). Furthermore, a meta-analysis of differentially expressed genes in patients with PD identified the downregulation of PSS1 in the SNpc (57), suggesting that although dysregulated PS is involved in pathogenesis, the cell may compensate for phospholipid defects through multiple mechanisms. As reported by us and others, data suggest that these PS alterations are deeply associated with αSyn pathology and neurological deficits (58–60), in agreement with previous observations (61–63). On the other hand, these data do not clarify whether these phospholipid alterations occur in all cell types in most affected brain areas or just in populations with higher levels of αSyn aggregates.

The consequences of these high levels of PS species bound to PUFAs in neurons and other brain cells are not well understood; however, they may include enhanced vulnerabilities to cell death mechanisms, including both cell-autonomous mechanisms such as ferroptosis (37) and non–cell-autonomous mechanisms. At the plasma membrane, the loss of PS symmetry and the presence of extracellular-facing PS on the plasma membrane have been identified in the brains of patients with both AD and mild cognitive impairment (64). Moreover, PS externalization is an “eat-me” signal for immune cells, including T cells, which designates the cell for immune attack and serves as an early indicator of apoptosis. In diseases associated with αSyn accumulation, the activation of microglia, T-cell ingress and release of pro-inflammatory cytokines are all observed at the respective sites of neurodegeneration, suggesting that αSyn defects contribute to PS via impacts on MAM regulation, predisposing vulnerable cells to immune attack and death during disease progression.

The MAM domains serve as the locus for sphingolipid enzymes. Therefore, the disruption of MAM by a pathologic dosage of αSyn can explain the unbalance in the levels of sphingolipid species towards a reduction in those containing long-fatty acyl chains and an increase in those bound to shorter FA. Notably, iPSC-derived αSyn-KO neurons showed an inverse phenotype from αSyn-Duplication neurons, directly implicating αSyn in the regulation of sphingolipid metabolism.

Defects in sphingolipid regulation, caused by mutations in *GBA1*, have also been linked to increases in αSyn aggregation (65). Altogether, these data reinforce the notion of a bidirectional interplay between αSyn regulation and sphingolipid homeostasis (66), which may explain why *GBA1* mutations in the context of Gaucher disease are associated with increased PD risk. These findings align with the existence of a bidirectional regulation between αSyn and sphingolipids (67), providing insights into the potential link between mutations in *GBA1* and *SNCA* that connect sphingolipid dyshomeostasis with increased PD risk. These sphingolipid alterations, however, were not statistically significant in striatum or cerebellar samples from MSA cases.

The modulation of enzymes involved in the synthesis and acylation of triglycerides, such as the acyl-CoA: diacylglycerol acyltransferase 2, also take place at the MAM (68). Interestingly, we identified higher levels of specific TG species exclusively in the ER of αSyn-Duplication. Considering our data, we posit that elevated TG levels in the ER could be a consequence of dysregulated MAM function and structure provoked by elevated αSyn. In turn, αSyn affinity for TG-rich membranes can explain the high levels of αSyn observed in ER fractions isolated from αSyn-Duplication neurons. The enrichment and aggregation of αSyn has also been associated with ER fragmentation and impaired function (69).

Among the different TGs species, those containing oleic acid and PUFAs were remarkably increased in αSyn-Duplication neurons, which agrees with the elevations in this type of TG species observed in sera from PD cases with the pathogenic αSyn-A53T mutation (69). Notably, other studies have reported that blocking the oleic acid–generating enzyme, stearoyl-CoA-desaturase, which is also located at the MAM domain, rescues αSyn-mediated toxicity in human neurons and mice carrying another structurally altered pathogenic point mutation (70, 71).

In summary, our study provides a novel framework for understanding the role of αSyn in lipid metabolism at MAM, the disruption of which may contribute to the development of PD. Moreover, our data clarify the source of the well-known lipid alterations in patients with PD associated with αSyn mutations. In addition, our approach could reveal new disease biomarkers that may be developed into a new tool for determining PD risk, improving the accuracy of PD diagnoses, and predicting PD progression, mainly if analyzed jointly with other PD hallmarks.

## Materials and Methods

### Cell Lines

The iPSC lines used in this study are shown in **Supplementary Table 4**; culture conditions have been previously described (32). The generation of the αSyn-KO line using the CRISPR-Cas9 system was previously described (30, 37).

### Subcellular Fractionation of IPSC-derived Neurons

A minimum of five confluent 15 cm^2^ plates of each cell line were subjected to directed differentiation. At DIV30, the plates were combined for each cell line and subjected to subcellular fractionation to obtain mitochondrial, ER, MAM, cytosolic, and crude mitochondrial (CM) fractionations, as previously described (72, 73). A minimum of four biological replicates were used per cell line.

### Protein Immunoblotting

Protein immunoblotting was performed as previously described (30) using 4%–20% Tris-Glycine gels (Invitrogen; XP04205) to separate 20 µg/µL of denatured protein. A dry transfer was performed for all samples using the iBlot2 (Invitrogen; IB21001), and blots were probed using antibodies recognizing dopamine transporter (Sigma-Aldrich; D6944), vesicular monoamine transporter 2 (Santa Cruz; sc-374079), monoamine oxidase A (Proteintech Group; 10539-1-ap), tyrosine hydroxylase (Millipore; AB152), αSyn (C-42) (BD Transduction Labs; 610786), β-actin (Sigma-Aldrich; A5441), Erp72 (Cell Signaling; D70D12), ATP5A1 (Invitrogen; 459240), ERLIN-2 (Cell Signaling; 2959S), protein kinase C (Sigma-Aldrich; P5704), ACSL4 (Abgent; AP2536b), PSS1 (Abcam; Ab157222), and PSS2 (Abcam; Ab183504). Membranes were washed in phosphate-buffered saline with 0.1% Tween 20 and probed with horseradish peroxidase-labeled anti-rabbit (Amersham; NA934V) or anti-mouse (Amersham; NA931V) secondary antibodies. The target bands were developed by enhanced chemiluminescence detection reagents (ThermoFisher; 34095 & 34580) and detected on the iBright 1500 Imaging System (ThermoFisher; A44114). Densitometry was performed using Fiji software (74), and protein quantities were normalized where stated.

### Phosphatidylserine Biosynthesis

*De novo* phosphatidylserine synthesis was performed on iPSC-derived neurons at DIV30 following directed differentiation, as previously described (11, 72).

### Phosphatidylserine Synthase Enzymatic Activities

A minimum of three confluent 15 cm^2^ plates per cell line were subjected to directed differentiation. At DIV30, plates were combined for each cell line and subjected to subcellular fractionation to obtain CM, ER, and cytosolic fractions, as previously described (72, 73). A minimum of three biological replicates were used per cell line. Each subcellular fraction was quantified using the BCA assay according to the manufacturer’s instructions, and 100 µM of protein per fraction was incubated with 1 µCi PE, L-a-1-palmitoyl-2-arachidonyl [arachidonyl-1-^14^C] (American Radiolabeled Chemicals (ARC); 0855-10 µCi) for 45 minutes. Lipids were then extracted using the modified Bligh and Dyer method (75), and thin-layer chromatography was performed as previously described (11, 72). Radiolabeled samples were read on a liquid scintillation counter (Perkin Elmer, Waltham, MA).

### Lipidomics

Lipidomics profiling was performed using UPLC–MS/MS (76, 77). Lipid extracts were prepared from cell lysates spiked with appropriate internal standards using a modified Bligh and Dyer method (75) and analyzed on a platform comprising Agilent 1260 Infinity HPLC integrated to Agilent 6490A QQQ mass spectrometer controlled by Masshunter v7.0 (Agilent Technologies, Santa Clara, CA). Glycerophospholipids and sphingolipids were separated with normal-phase HPLC as described previously (78), with a few modifications. An Agilent Zorbax Rx-Sil column (2.1 × 100 mm, 1.8 µm) maintained at 25°C was used under the following conditions: mobile phase A (chloroform: methanol: ammonium hydroxide, 89.9:10:0.1, v/v) and mobile phase B (chloroform: methanol: water: ammonium hydroxide, 55:39:5.9:0.1, v/v); 95% A for 2 min, decreased linearly to 30% A over 18 min, and further decreased to 25% A over 3 min, before returning to 95% over 2 min and held for 6 min. Separation of sterols and glycerolipids was carried out on a reverse phase Agilent Zorbax Eclipse XDB-C18 column (4.6 × 100 mm, 3.5 µm) using an isocratic mobile phase, chloroform, methanol, 0.1 M ammonium acetate (25:25:1) at a flow rate of 300 µl/min.

Quantification of lipid species was accomplished using multiple reaction monitoring transitions (78–80) under both positive and negative ionization modes in conjunction with referencing of appropriate internal standards (lipid species abbreviations can be found in **Supplementary Table 3**): PA 14:0/14:0, PC 14:0/14:0, PE 14:0/14:0, PG 15:0/15:0, PI 17:0/20:4, PS 14:0/14:0, BMP 14:0/14:0, APG 14:0/14:0, LPC 17:0, LPE 14:0, LPI 13:0, Cer d18:1/17:0, SM d18:1/12:0, dhSM d18:0/12:0, GalCer d18:1/12:0, GluCer d18:1/12:0, LacCer d18:1/12:0, D7-cholesterol, CE 17:0, MG 17:0, 4ME 16:0 diether DG, D5-TG 16:0/18:0/16:0 (Avanti Polar Lipids, Alabaster, AL). Lipid levels for each sample were calculated by summing up the total number of moles of all lipid species measured by all three LC-MS methodologies and then normalizing the total to mol %. The final data are presented as mean mol % with error bars representing standard deviation (SD). RF classification was performed as previously described (13, 16).

### Statistical Analysis

Data are presented as the mean ± SD. All averages are the result of three or more independent experiments. iPSC-derived neurons from the same individual were differentiated at different times from different NPC passages. The data distribution was assumed to be normal with the differences between means analyzed using either one-way or two-way repeated measures analysis of variance (ANOVA), followed by Tukey’s multiple comparison post hoc test, as reported in each figure legend. GraphPad Prism (Version 10, GraphPad Software Inc., USA) was used for statistical analyses. In all analyses, the null hypothesis was rejected at the 0.05 level. The investigators were not blinded when quantifying imaging experiments. *p<0.05, **p<0.01, ***p<0.001, ****p<0.0001.

### Study Approval

An exempt protocol was approved by the Institutional Review Board at Columbia University Irving Medical School for de-identified human pathologic specimens.

## Data, Materials, and Software Availability

All study data are included in the article and/or SI Appendix.

## Acknowledgments

We would like to acknowledge and thank Renu Nandakumar, Yimeng (Oliver) Xu, and Laura DeFreitas of the Biomarkers Core (Columbia University) for extracting and processing the lipidomic samples. We would also like to acknowledge Benoit Giasson (University of Florida) for kindly providing the αSyn antibodies 3H11, 94-3A10, and 1D12. This research was supported by the William N. and Bernice E. Bumpus Foundation (P.B is a recipient of the Early Career Investigator Innovation Award (WBBF CU22-0241), plus grants from the Fond National de Recherche within the INTER program (INTER/LEIR/18/12719318) to PB and RK, and PEARL (FNR/P13/6682797) to RK, and the National Centre for Excellence in Research on Parkinson’s disease (NCER-PD/11264123) program and by the European Union’s Horizon 2020 research and innovation program under Grant Agreement No 692320 (WIDESPREAD; CENTRE-PD) to RK. This work was further supported by the US National Institutes of Health (NS107442, NS117583, NS111176, AG064596) to SP, (NS121826, NS133979) to UD, and (AG056387) to EAG, the DoD (W81XWH-22-1-0127) and the Parkinson Foundation to SP, the Ludwig Family Foundation (GT006761) to SP and PB, the Michael J Fox Foundation to CG-L, the Leir Foundation (GT006967) and Art2Cure to PB, CG-L, RK and SP. The funders had no role in the design of the study, the collection, analyses, or interpretation of data, the writing of the manuscript, or the decision to publish the results.

## Author Contributions

PB, CG-L, ZKC, EB, EAG, and SP designed research; PB, CG-L, TY, and EAG performed research; PB, EAG, TY, MV, EAG, and SP analyzed data; provide and validate critical reagent XF, BFRS, SBL, NH, HR, AFT, and RK; and PB, NH, EB, UD, SF, RK, EAG, and SP wrote the paper.

## Supplemental Information

Supplementary Tables 1-4. Supplementary Figures 1-6

