## Supplementary Figures for "The Role of Alpha-Synuclein in Synucleinopathy: Impact on Lipid Regulation at Mitochondria–ER Membranes"

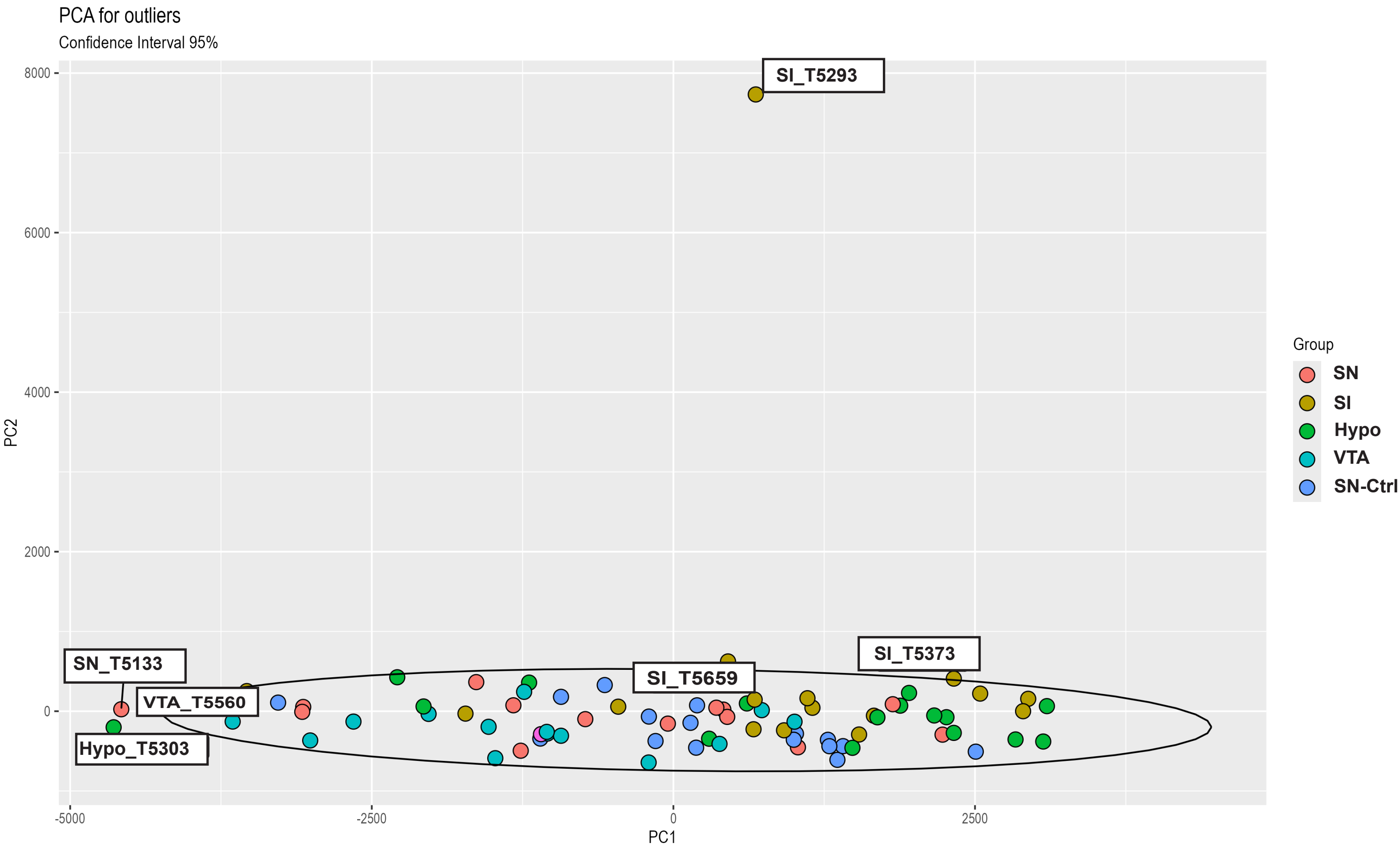


**Supplementary Fig.1. Principle Component Analysis (PCA) of postmortem brain regions.** Sam-ples outside the 95% confidence interval (red eliptical box) were excluded from this study. Abbrevia-tions: substantia nigra (SN), substantia innominata (SI), hypothalamus (Hypo), ventral tegmental area (VTA), SN from non PD donor controls (SN-Ctrl).

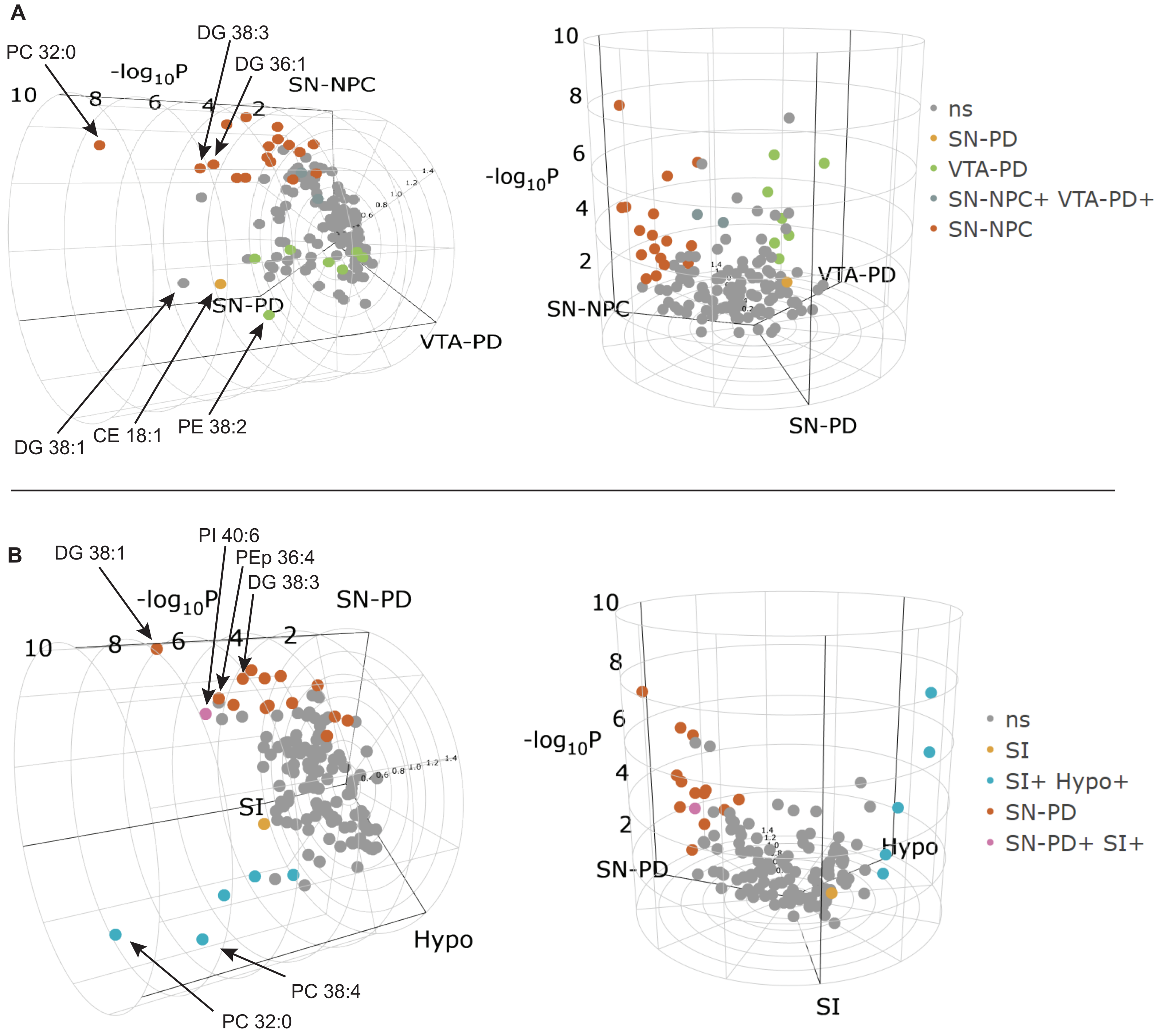


**Supplementary Fig.2A. Cylindrical volcano plots of significantly altered lipids from different brain regions.** 3D volcano plots were assembled using the p values calculated from a one-way ANOVA, followed by pairwise t test corrected by false discovery rate (FDR) of 5 tissues: substantia nigra (SN), ventral tegmental area (VTA), hypothalamus (Hypo), substantia innominata (SI), all from PD donors, and the substantia nigra from non PD control donors (NPC). The Z axis of the volcano plot shows ANOVA p values (-log10) for the associated lipid species for the subset of 3 groups as illustrated in: **(A)** SN-PD, VTA-PD, SN-NPC; and **(B)** SN-PD, SI, Hypo. The radial axis corresponds to the Z-score. The data for 3 subset groups were reduced to a 2D polar coordinate system. Colored dots indicate which species are altered significantly by adjusted p values in pairwise comparisons.

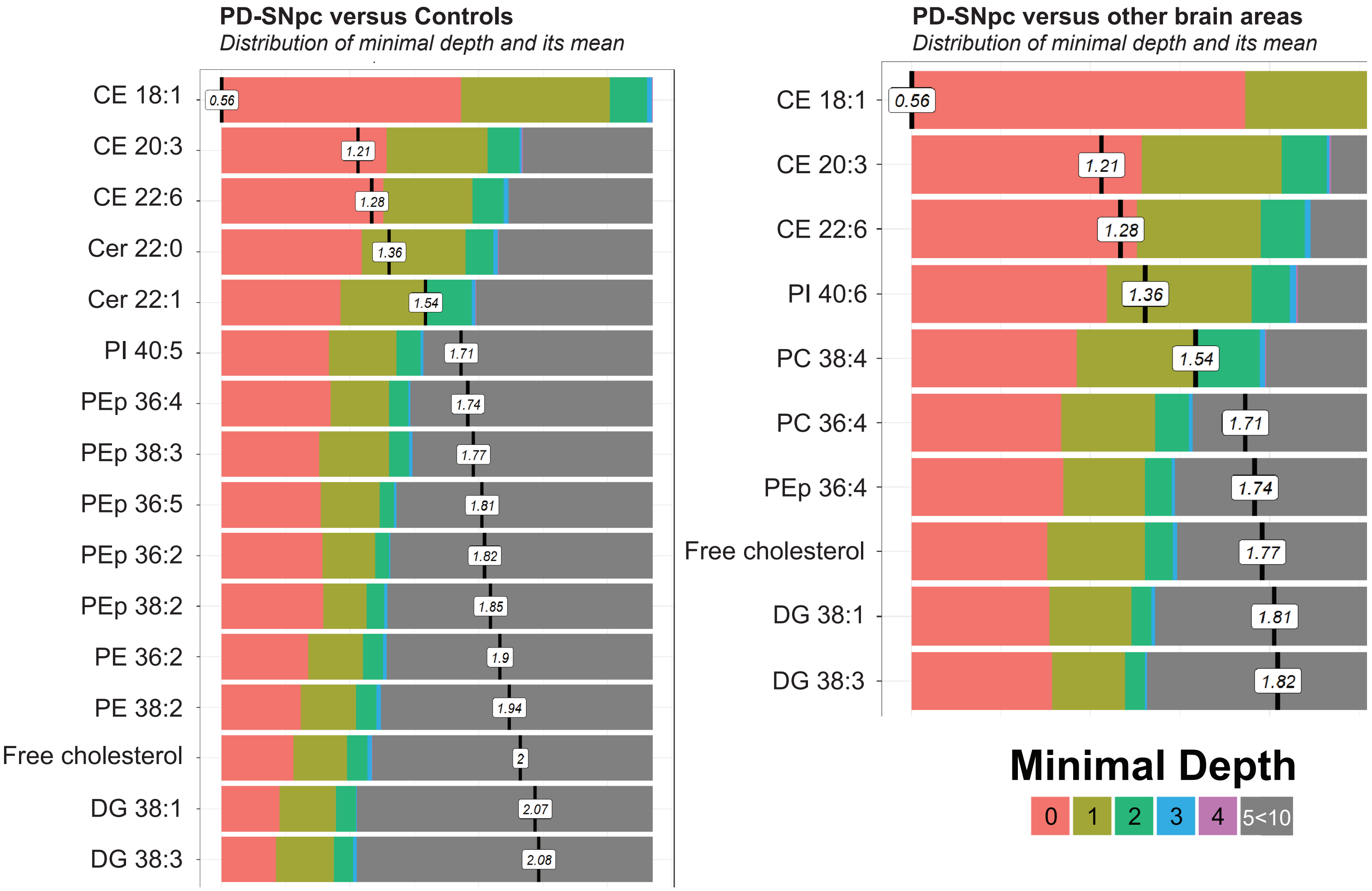


**Supplementary Fig.2B. Variable importance based on minimum depth from the Random Forest (RF) analysis.** Note, minimal depth indicates how early a lipid is involved in decision trees. Higher frequencies at lower nodes indicate that specific lipid species effectively classify the different groups.

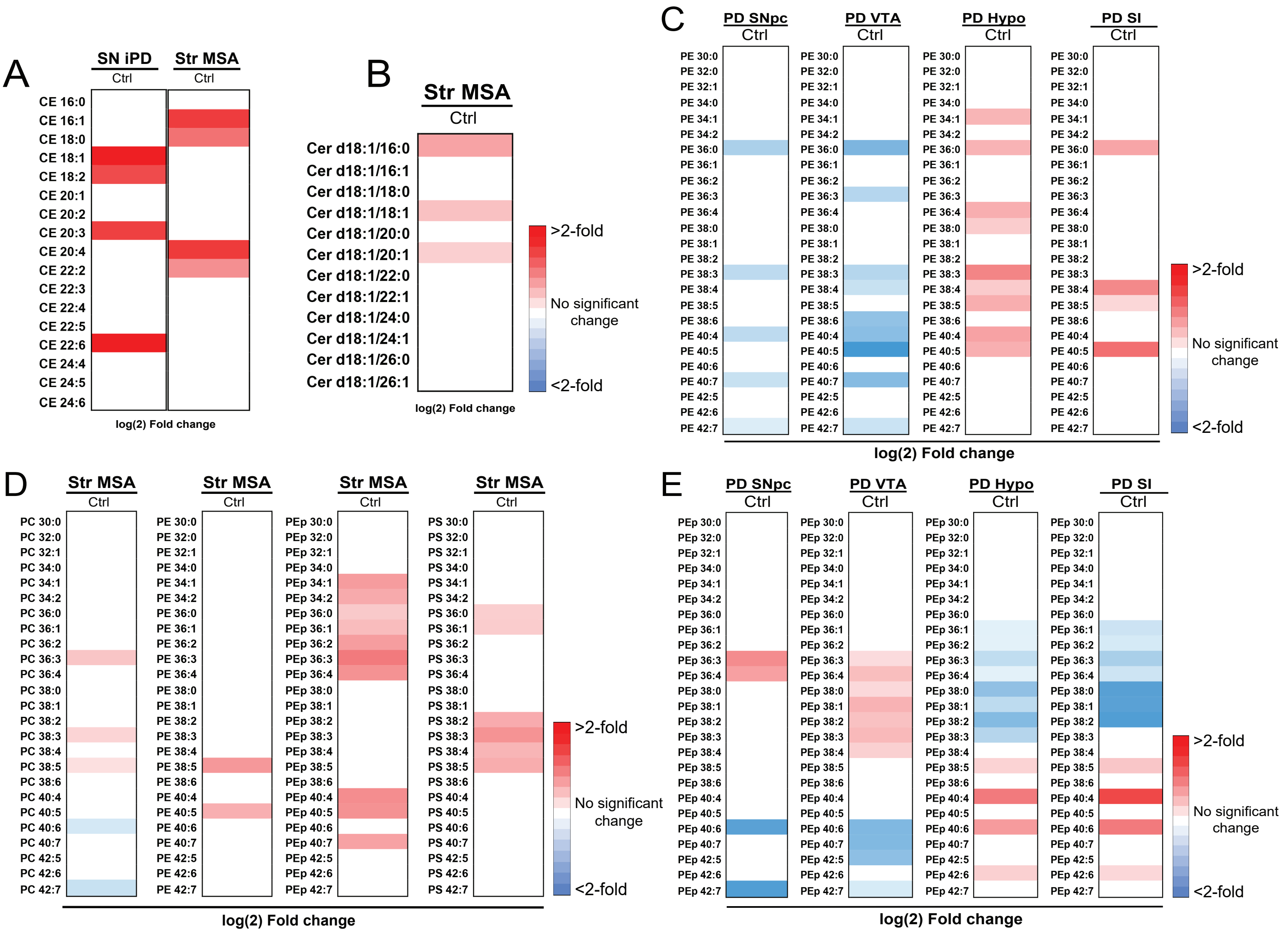


**Supplementary Fig.3. Lipid alterations in postmortem brains.** Lipidomic heat map showing Log(2) fold changes in different brain regions of postmortem Parkinson's disease (PD) and multiple system atrophy (MSA) donors. **(A)** Fold-increases in the levels of specific cholesterol esters (CE) in the substantia nigra pars compacta (SN) of PD donors and striatum (Str) of MSA donors compared to Controls. **(B)** Increases in the levels of ceramide (Cer) species in the striatum of MSA donors compared to Controls. **(C)** Alterations of phosphatidylethanolamine (PE) lipid species in different regions of PD brain (substantia nigra pars compacta (SNpc), ventral tegmental area (VTA), hypothalamus (Hypo), substantia innominata (SI)) at postmortem. **(D)** Alterations in different species of phospholipids (phosphatidylcholine (PC), PE, plasmalogen phosphatidylthanolamine (PEp) and phosphatidylserine (PS)) in the striatum of MSA donors compared to controls. **(E)** Alterations in different species of PEp in different regions of PD brains at postmortem compared to controls.

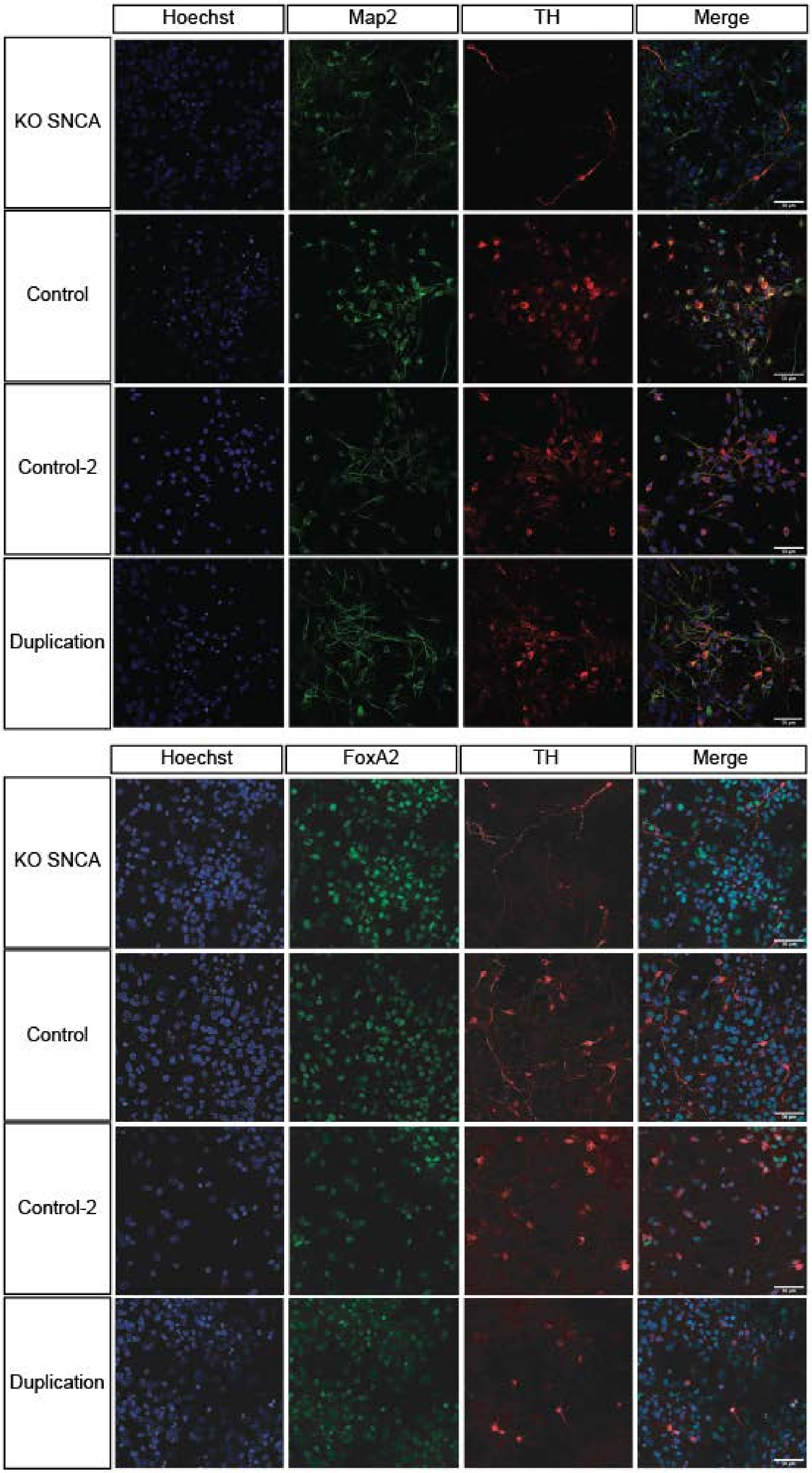


**Supplementary Fig.4A. Characterization of midbrain dopaminergic neurons**. Representative images of neurons fluorescently labelled with the mature neuronal marker Map2, the rate-limiting enzyme for catecholamine biosynthesis and the biosynthesis of dopamine tyrosine hydroxylase (TH), and the ventral midbrain marker FoxA2.

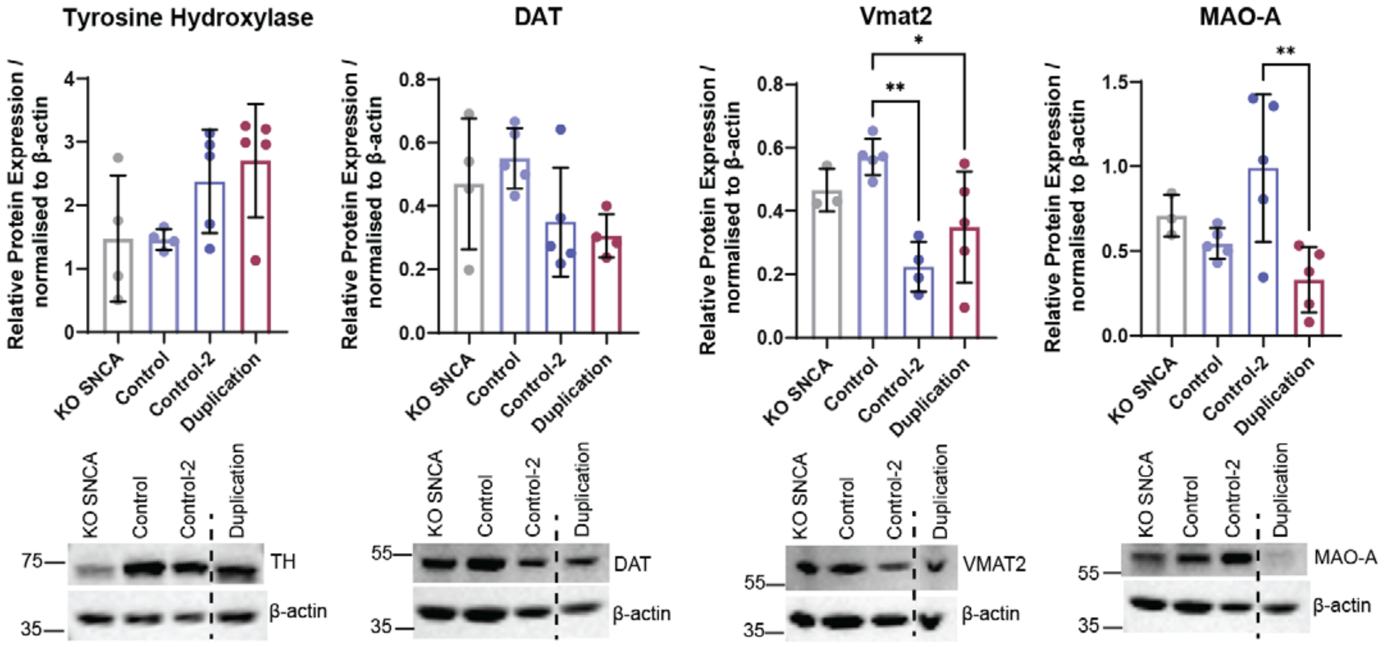


**Supplementary Fig.4B. Neuronal chacterization.** Cell lines were positive for markers of dopaminergic neurons as shown by a representative western blot. Data are means ± SD of a minimum of 3 independent biological replicates (n) analyzed by an ordinary one-way ANOVA, for statistical analyses of the indicated proteins a Tukey's multiple comparison post-hoc was used with a single pooled variance. Tyrosine Hydroxylase (TH) (Interaction $F_{3,14}$ =2.821; p=0.0772), DAT (Interaction $F_{3,14}$ =2.706; p=0.0852), VMAT2 (Interaction $F_{3,13}$ =7.841; **p=0.0031), MAO-A (Interaction $F_{3,14}$ =5.522; *p=0.0103). In all statistical analyses: *p<0.05, **p<0.01, ***p<0.001, ****p<0.0001.

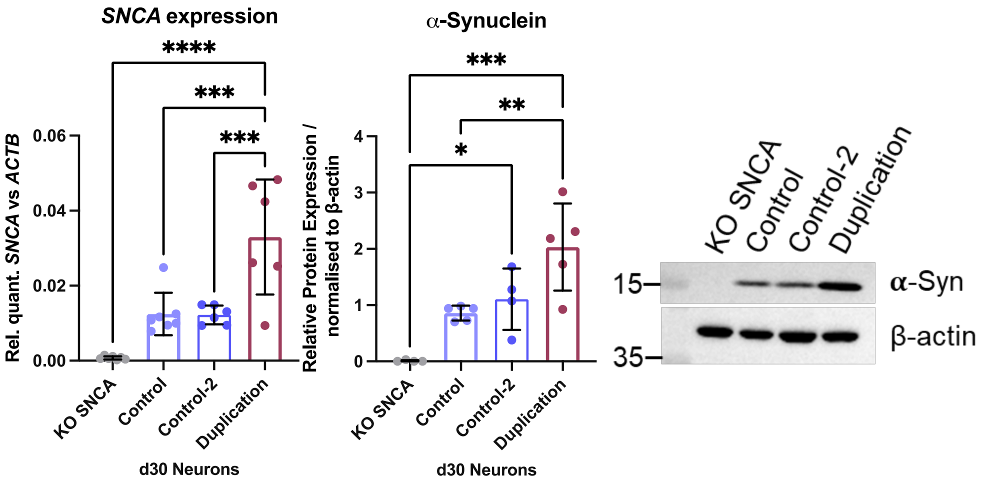


**Supplementary Fig. 4D. Validation of iPS-derived neurons expressing SNCA.** Gene and protein expression of αSyn after 30-days of directed neuronal differentiation. For SNCA gene expression the data are means ± SD of a minimum of 6 independent biological replicates (n) analyzed by an ordinary one-way ANOVA with (A) Interaction $F_{3,22}$ =17.98; ****p<0.0001. Characterization of αSyn protein level by western blot, the data was obtained using the means ± SD of a minimum of 4 independent biological replicates (n) analyzed by an ordinary one-way ANOVA (Interaction $F_{3,14}$ =13.04; ***p=0.0002) with a representative image shown. For all statistical analysis, a Tukey's multiple comparison post-hoc was used with a single pooled variance.

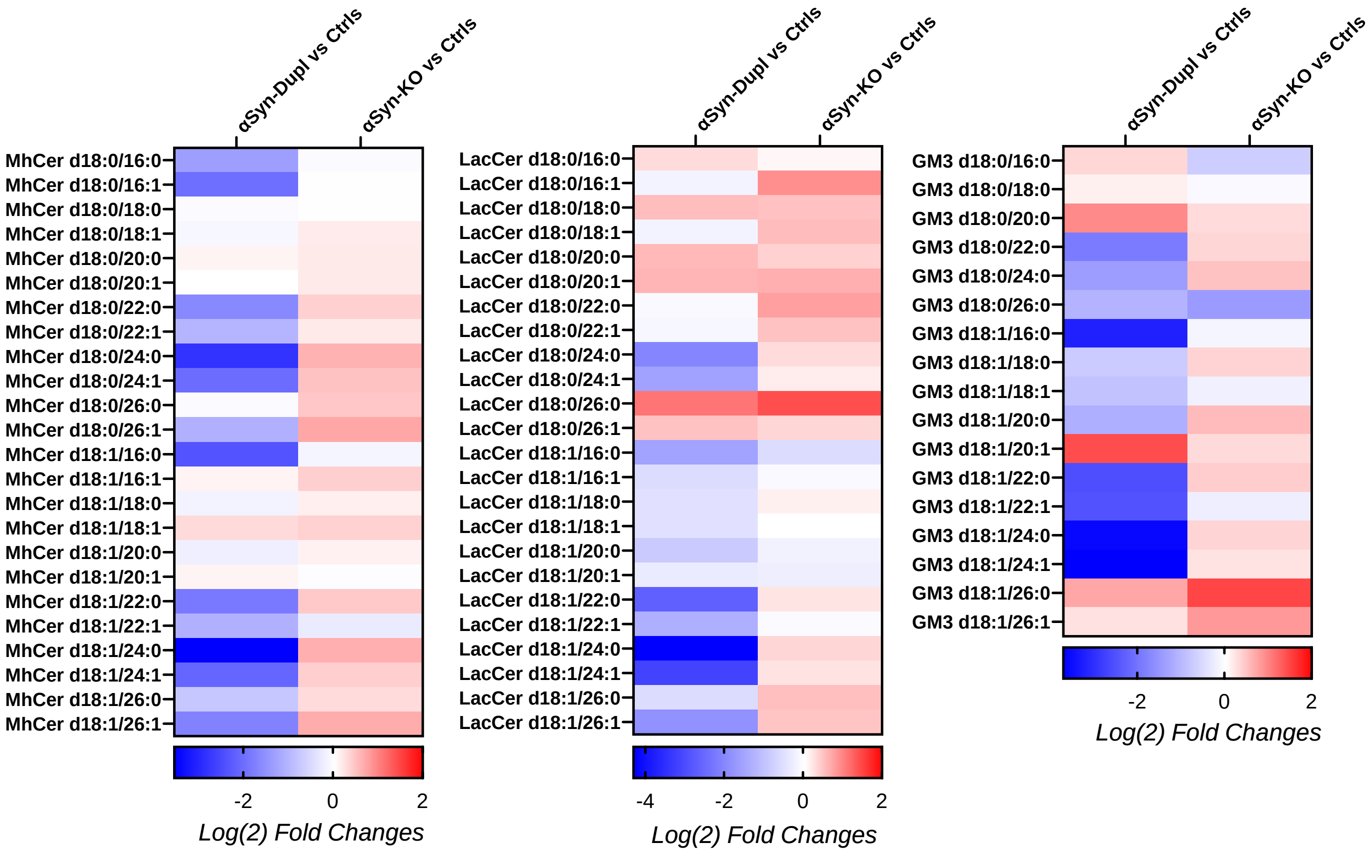


**Supplementary Fig.4E. Altered complex sphingolipid and glycolipid classes in patient-derived neurons expressing differing αSyn dosage.** Lipidomic heat map of Log(2) fold changes of selected lipid species in iPS-derived neurons carrying altered αSyn dosage compared to Controls. Lipid abbreviations: MhCer: Monohexosylceramide; LacCer: Lactosylceramide; GM3: Monosialodihexosylganglioside.

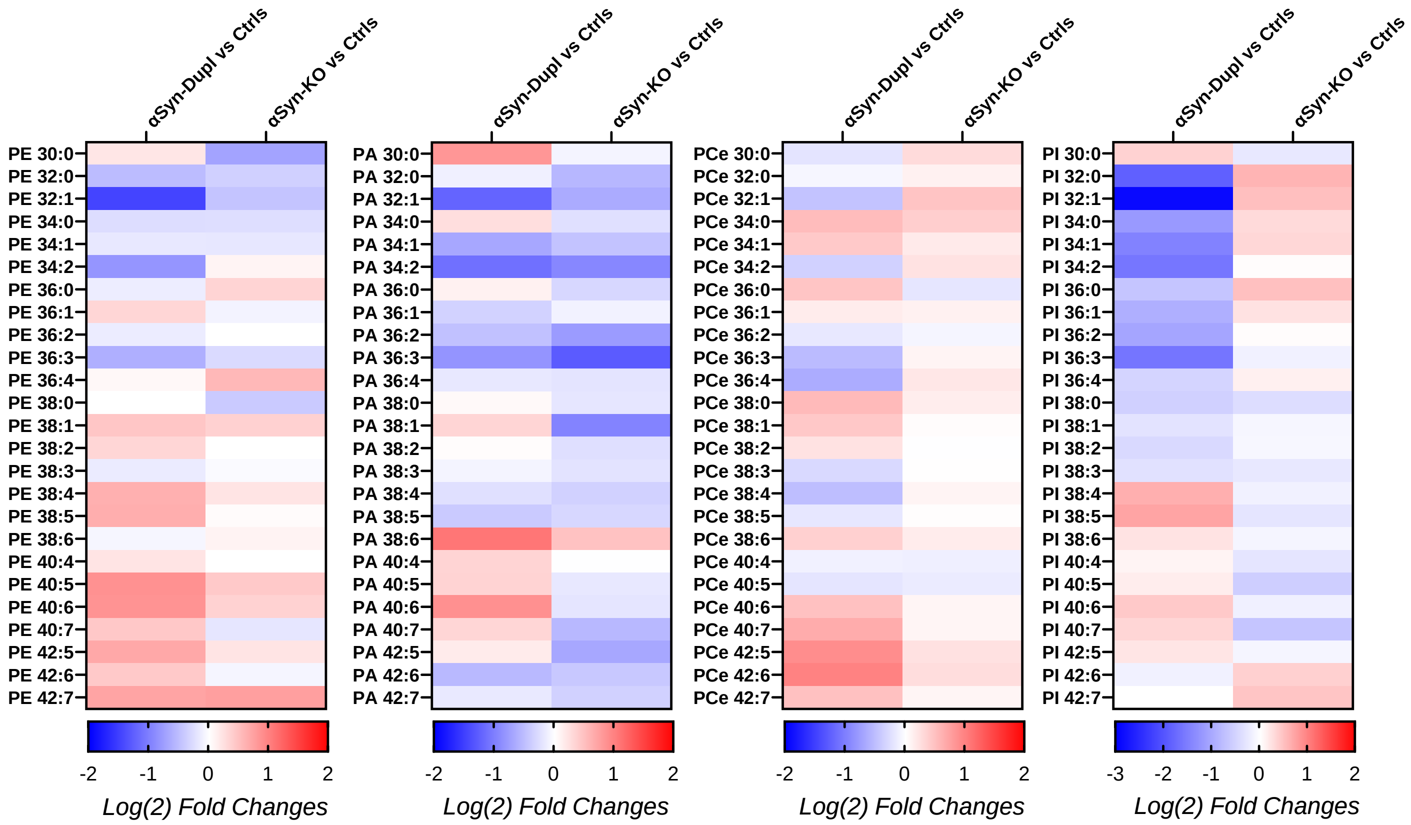


**Supplementary Fig. 4F: Altered phospholipid classes in neurons expressing differing αSyn dosage.** Lipidomic heat map of Log(2) fold changes of selected lipid species in iPS-derived neurons carrying altered αSyn dosage compared to Controls. Lipid abbreviations: PE: Phosphatidylethanolamine; PA: Phosphatidic Acid; PCe: Ether phosphatidylcholine; PI: Phosphatidylinositol.

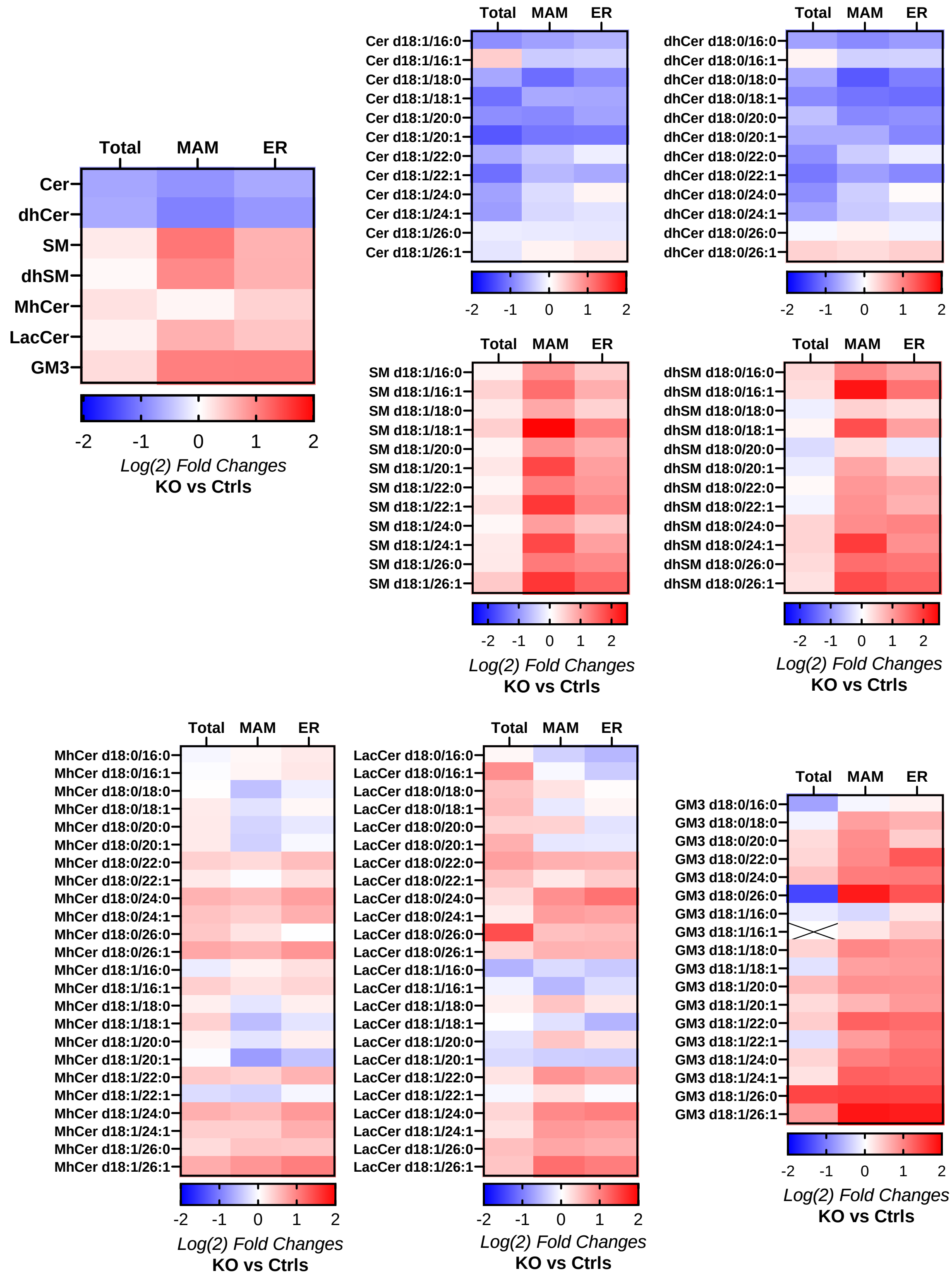


**Supplementary Fig.5A. Lipid alterations in αSyn-KO neurons across different subcellular fractions.** Lipidomic heat map of Log(2) fold changes of sphingolipid and glycolipid lipid classes and lipid species in iPS-derived neurons carrying αSyn-KO compared to Controls in the total non-fractionated homogenate, MAM and bulk-ER. Lipid abbreviations: Cer: Ceramide; dhCer: Dihydroceramide; SM: Sphingomyelin; dhSM: Dihydrosphingomyelin; MhCer: Monohexosylceramide; LacCer: Lactosylceramide; GM3: Monosialodihexosylganglioside.

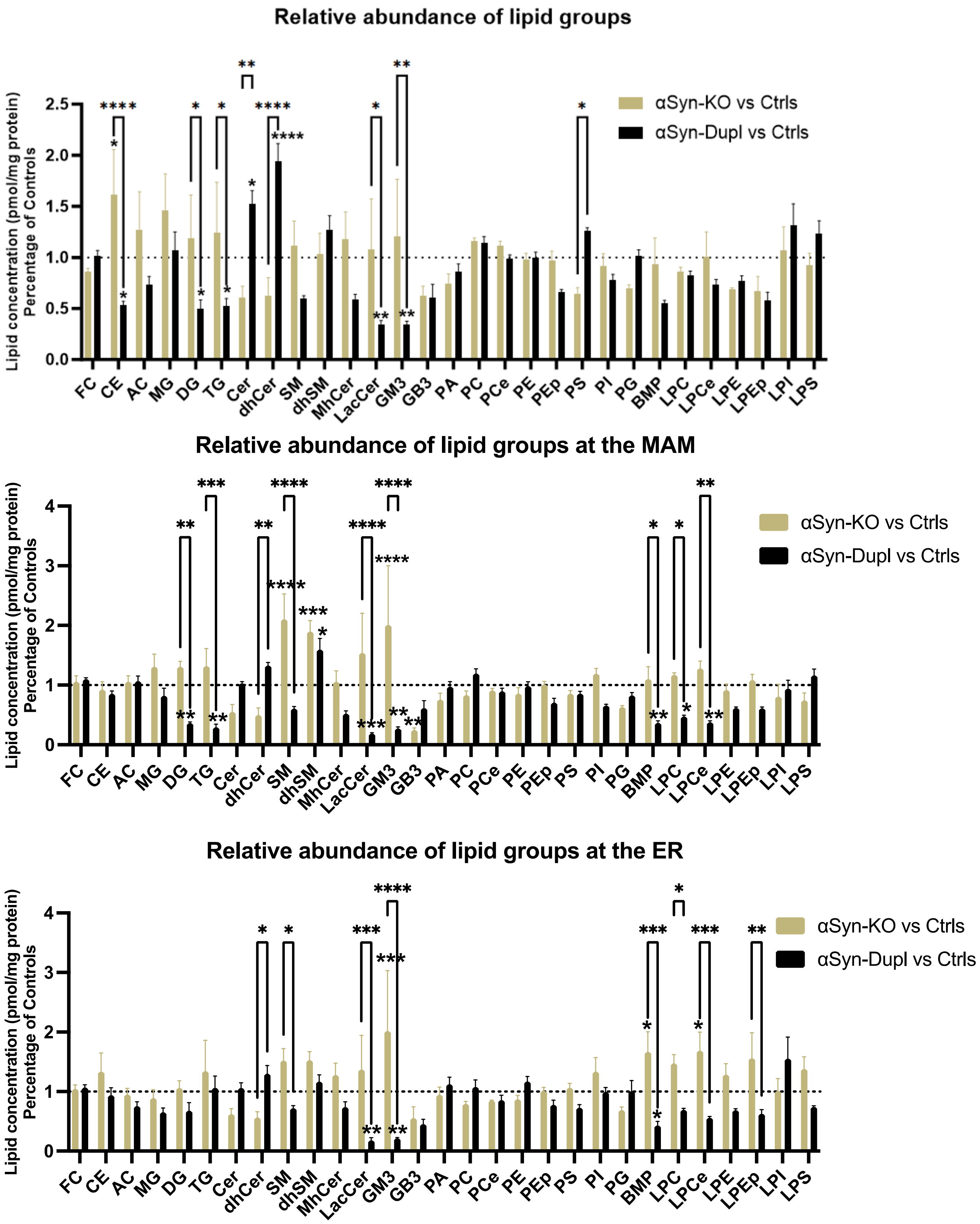

Relative abundance of lipid groups
Lipid concentration (pmol/mg protein) Percentage of Controls
αSyn-KO vs Ctrls
αSyn-Dupl vs Ctrls
FC CE AC MG DG TG Cer dhCer SM dhSM MhCer LacCer GM3 GB3 PA PC PCe PE PEp PS PI PG BMP LPC LPCe LPE LPEp LPI LPS
Relative abundance of lipid groups at the MAM
Lipid concentration (pmol/mg protein) Percentage of Controls
αSyn-KO vs Ctrls
αSyn-Dupl vs Ctrls
FC CE AC MG DG TG Cer dhCer SM dhSM MhCer LacCer GM3 GB3 PA PC PCe PE PEp PS PI PG BMP LPC LPCe LPE LPEp LPI LPS
Relative abundance of lipid groups at the ER
Lipid concentration (pmol/mg protein) Percentage of Controls
αSyn-KO vs Ctrls
αSyn-Dupl vs Ctrls
FC CE AC MG DG TG Cer dhCer SM dhSM MhCer LacCer GM3 GB3 PA PC PCe PE PEp PS PI PG BMP LPC LPCe LPE LPEp LPI LPS

**Supplementary Fig.5B. Relative abundance of lipid groups expressed in patient-derived neurons differing αSyn dosage in different cellular subcellular compartments.** Relative lipid concentration (pmol/mg protein) normalized to controls. Data are means ± SEM of a minimum of 3 independent biological replicates (n) analyzed by an ordinary two-way ANOVA. Post-hoc analysis was performed using Tukey's multiple comparison test, for the relative abundance of lipid groups at the Total homogenate: Interaction ($F_{56,348}$ = 2.802; ****p<0.0001) and Column Factor (cell line) ($F_{2,348}$ = 5.848; **p=0.0032); the MAM: Interaction ($F_{56,348}$ = 3.533; ****p<0.0001), and Column Factor (cell line) ($F_{2,348}$ = 23.37; ****p<0.0001), and at the ER: Interaction ($F_{56,348}$ = 2.257; ****p<0.0001), and Column Factor (cell line) ($F_{2,348}$ = 18.17; ****p<0.0001). Lipid abbreviations: FC: Free Cholesterol; CE: Cholesterol Ester; AC: Acyl Carnitine; MG: Monoacylglycerol; DG: Diacylglycerol; TG: Triacylglycerol; Cer: Ceramide; dhCer: Dihydroceramide; SM: Sphingomyelin; dhSM: Dihydrosphingomyelin; MhCer: Monohexosylceramide; LacCer: Lactosylceramide; GM3: Monosialodihexosylganglioside; GB3: Globotriaosylceramide; PA: Phosphatidic Acid; PC: Phosphatidylcholine; PCe: Ether phosphatidylcholine; PE: Phosphatidylethanolamine; PEp: Plasmalogen phosphatidylethanolamine; PS: Phosphatidylserine; PI: Phosphatidylinositol; PG: Phosphatidylglycerol; BMP: Bis(monoacylglycerol)phosphate; LPC: Lysophosphatidylcholine; LPCe: Ether Lysophosphatidylcholine; LPE: Lysophosphatidylethanolamine; LPEp: Plasmalogen Lysophosphatidylethanolamine; LPI: Lysophosphatidylinositol; LPS: Lysophosphatidylserine.

### Subcellular fractions from αSyn-KO iPS-derived neurons vs Controls

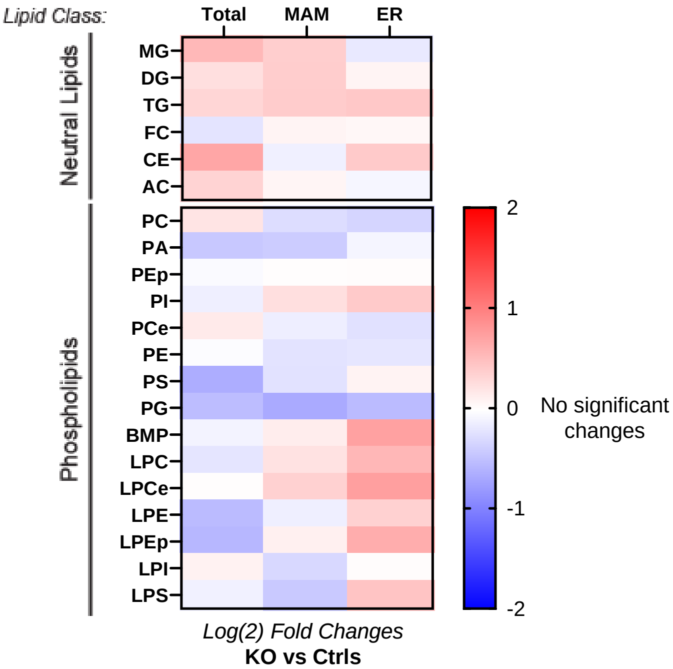


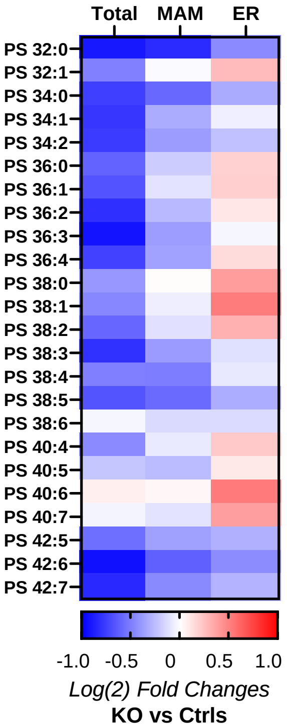


**Supplementary Fig.5C. Lipid alterations in αSyn-KO neurons across different subcellular fractions.** Lipidomic heat map of Log(2) fold changes of neutral lipid and phospholipid lipid classes and PS lipid species in iPS-derived neurons carrying αSyn-KO compared to Controls in the total non-fractionated homogenate, MAM and bulk-ER. Lipid abbreviations: MG: Monoacylglycerol; DG: Diacylglycerol; TG: Triacylglycerol; FC: Free Cholesterol; CE: Cholesterol Ester; AC: Acyl Carnitine; PC: Phosphatidylcholine; PA: Phosphatidic Acid; PEp: Plasmalogen phosphatidylethanolamine; PI: Phosphatidylinositol; PCe: Ether phosphatidylcholine; PE: Phosphatidylethanolamine;; PS: Phosphatidylserine; PG: Phosphatidylglycerol; BMP: Bis(monoacylglycerol)phosphate; LPC: Lysophosphatidylcholine; LPCe: Ether Lysophosphatidylcholine; LPE: Lysophosphatidylethanolamine; LPEp: Plasmalogen Lysophosphatidylethanolamine; LPI: Lysophosphatidylinositol; LPS: Lysophosphatidylserine

### Crude Mitochondrial Fraction

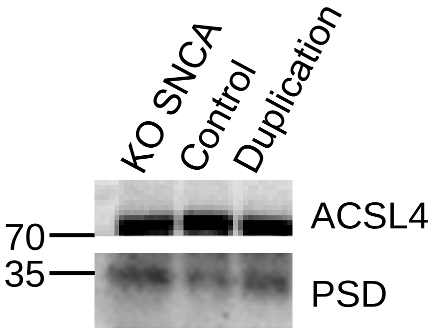


**Supplementary Fig.6. Expression of PSD in iPS-derived neurons in a crude mitochondrial subcellular fraction**
