## Supplementary Tables for "The Role of Alpha-Synuclein in Synucleinopathy: Impact on Lipid Regulation at Mitochondria–ER Membranes"

| **Supplementary Table 1. Substantia nigra sample donors' demographic Information** | | | | |
|---|---|---|---|---|
| **Group** | **Sample size** | **Age (Year ± SD)** | **Sex (M/F)** | **PMI (hour ± SD)** |
| Control | 14 | 80.2 ± 12.3 | 8/6 | 14.1 ± 7.5 |
| PD | 16 | 79.7 ± 7.1 | 12/4 | 18.6 ± 12.1 |

Abbreviations: PD, Parkinson's disease; PMI, post-mortem interval; SD, standard deviation; M, male; F, female.

Supplementary Table 2: Lipids detected (>500 species belonging to 31 classes).

| Abbr. | Lipid (detectable species) | Abbr. | Lipid (detectable species) |
|---|---|---|---|
| **FC** | Free Cholesterol | **PC** | Phosphatylcholine (25 species) |
| **CE** | Cholesterol Ester (20 species) | **PCe** | Ether phosphatidylcholine (25 species) |
| **AC** | Acyl Carnitine (9 species) | **PE** | Phosphatidylethanolamine (25 species) |
| **MG** | Monoacylglycerol (18 species) | **PEp** | Plasmalogen phosphatidylethanolamine (25 species) |
| **DG** | Diacylglycerol (28 species) | **PS** | Phosphatidylserine (25 species) |
| **TG** | Triacylglycerol (42 species) | **PI** | Phosphatidylinositol (25 species) |
| **dhCer** | Dihydroceramide (12 species) | **PG** | Phosphatidylglycerol (25 species) |
| **Cer** | Ceramide (12 species) | **BMP** | Bis(monoacylglycero)phosphate (25 species) |
| **SM** | Sphingomyelin (12 species) | **AcylPG** | Acyl Phosphatidylglycerol (15 species) |
| **dhSM** | Dihydrosphingomyelin (12 species) | **LPC** | Lysophosphatidylcholine (9 species) |
| **Sulf** | Sulfatide (18 species) | **LPCe** | Ether lysophosphatidylcholine (9 species) |
| **MHCer** | Monohexosylceramide (24 species) | **LPE** | Lysophosphatidylethanolamine (9 species) |
| **LacCer** | Lactosylceramide (24 species) | **LPEp** | Plasmogen Lysophosphatidylethanolamine (9 species) |
| **GM3** | Monosialodihexosylganglioside (18 species) | **LPI** | Lysophosphatidylinositol (9 species) |
| **GB3** | Globotriaosylceramide (12 species) | **LPS** | Lysophosphatidylserine (11 species) |
| **PA** | Phosphatidic acid (25 species) | | |

Abbreviations: Abbr., abbreviation.

**Supplementary Table 3. Striatum sample donors' demographic Information**

| Group | Sample size | Age (Year ± SD) | Sex (M/F) | PMI (hour ± SD) |
|---|---|---|---|---|
| Control | 16 | 68.2 ± 7.9 | 13/3 | 13.5 ± 7.9 |
| PD/DLB | 11 | 77.9 ± 5.9 | 9/2 | 16.2 ± 10.9 |
| MSA | 10 | 68.6 ± 8.5 | 5/5 | 15.9 ± 9.7 |

Abbreviations: PD, Parkinson's disease; DLB, dementia with Lewy bodies; MSA, multiple system atrophy; PMI, post-mortem interval; SD, standard deviation; M, male; F, female.

Supplementary Table 4: List of cell lines

| **Cell name** | **Fibroblast name (iPSC clone)** | **Status** | **Sex (M/F)** | **Age at Biopsy** | **iPSC reprogramming method** | **Mutation** | **Reference** |
|---|---|---|---|---|---|---|---|
| KO SNCA | 17608 (c.6) (Gene-edited c.21) | Control with no αSyn | M | 67 | - | Deletion in Exon 2 leads to loss of αSyn | (1, 2) |
| Control | 16426 (c.33) | Control | M | 72 | Sendai | - | (3) |
| Control-2 | 18075 (c.5) | Control | F | 72 | Sendai | - | (4) |
| Duplication | Dup. (A13) | PD | F | 67 | Sendai | SNCA gene locus duplication | (5) |

Abbreviations: iPSC, induced pluripotent stem cell; M, male; F, female; KO, knock out; Dup., duplication; PD, Parkinson's disease; αSyn, alpha-synuclein.
